# Streamlining effects of extra telomeric repeat on telomere folding revealed by fluorescence-force spectroscopy

**DOI:** 10.1101/651919

**Authors:** Jaba Mitra, Taekjip Ha

## Abstract

Tandem repeats of guanine rich sequences are ubiquitous in the eukaryotic genome. For example, in the human cells, telomeres at the chromosomal ends comprise of kilobases repeats of T_2_AG_3_. Four such repeats can form G-quadruplexes (GQs). Biophysical studies have shown that GQs formed from four consecutive repeats possess high diversity both in their structure and in their response to tension. In principle, a GQ can form from any four repeats that may not even be consecutive. In order to investigate the dynamics of GQ possessing such positional multiplicity, we studied five and six repeats human telomeric sequence using single molecule FRET as well as its combination with optical tweezers. Our results suggest preferential formation of GQs at the 3’ end both in K^+^ and Na^+^ solutions although minority populations with a 5’ GQ or long-loop GQs were also observed. Using a vectorial folding assay which mimics the directional nature of telomere extension, we found that the 3’ preference holds even when folding is allowed to begin from the 5’ side. Interestingly, the unassociated T_2_AG_3_ segment has a streamlining effect in that one or two mechanically distinct species was observed at a single position instead of six or more observed without an unassociated repeat. Location of GQ on a long G-rich telomeric overhang and reduction in diversity of GQ conformations and mechanical responses through adjacent sequences have important implications in processes such as telomerase inhibition, alternative lengthening of telomeres, T-loop formation, telomere end protection and replication.

## Introduction

As early as 1962, guanosine moieties were known to self-assemble into tetrameric structures via Hoogsteen hydrogen bonds (1). Planar association between four guanines by eight hydrogen bonds in the presence of a central coordinating monovalent cation (e.g. Na^+^ and K^+^) generates a G-quartet or a G-tetrad (2,3). Stacking of two or more G-quartets creates a very stable DNA secondary structure called the G-Quadruplex (GQ) (4). Complete sequencing of human genome has identified about 300,000 putative sequences that can fold into GQs (5). Expansion of GQ-forming motifs have been implicated in pathogenicity associated with several human neurological disorders (6). Additionally, GQ-forming sequences have been reported in various viral genomes including human immuno-deficiency virus, Epstein-Barr virus and human papillomavirus (7).

In human cells, telomeric DNA continues beyond the double-stranded region as a 3’ 100-200 nucleotides (nt) single-stranded DNA made of repeats of hexanucleotide T_2_AG_3_ (8). This G-rich overhang of T_2_AG_3_ repeats form GQ structures under physiological conditions (4,9,10). Telomere length homeostasis has important implications in cell survival and proliferation (11–14) and telomeric GQs are studied as targets for cancer treatment and for potential applications in nanotechnology (15–17).

Despite an apparently simple, repeating G-rich sequence, structural studies with circular dichroism (CD), nuclear magnetic resonance (NMR) and X-ray crystallography have revealed extreme polymorphism in human telomeric GQs composed of four T_2_AG_3_ repeats (18–24). Such diversity suggests co-existence of different conformations in the telomere with implications on rational designing of GQ-targeting drugs. GQ polymorphism and conformational dynamics in longer telomeric DNA containing five or more T_2_AG_3_ repeats, which should better mimic a telomere overhang, have been examined only in a few studies (25–29).

GQ formation requires a minimum of four G-rich segments (30). As human telomeric DNA contains multiple hexanucleotide (T_2_AG_3_) repeats and stretches for several kilobases, it has been assumed that GQs can form anywhere along the G-rich strand (31). Hence, under physiological conditions, a long telomeric DNA with more than four T_2_AG_3_ repeats can harbor different forms of terminal structures depending on the position of GQ(s). GQ locations on telomeric overhangs can modulate telomerase extension or mechanism of alternative lengthening of telomere (ALT) (14). An exonuclease hydrolysis assay suggested preferential GQ formation via association of four consecutive hexanucleotide repeats at the 3’ end of long telomeric DNA, leaving out a T_2_AG_3_ repeat at the 5’ end (26). Another study employing site-specific labeling of guanine residues suggested the presence of hybrid GQ structures, incorporating one or more hexanucleotide repeat within a single loop to connect adjacent Gs in a G-quartet (29). Ensemble average techniques used in the above studies are limited by time resolution and hence cannot univocally deconvolute individual species from a heterogeneous population. Dynamic exchange between GQ conformations in telomeric DNA spanning five to seven T_2_AG_3_ repeats have been observed using single molecule fluorescence resonance energy transfer (smFRET) (32) and single molecule optical tweezers studies of human telomeric DNA of four to seven repeats indicated that GQs more frequently form using four consecutive repeats (25). In a recent study we demonstrated extreme mechanical diversity heterogeneity of GQs in 22 nt long human telomeric DNA (four telomeric repeats, where at least six different mechanically distinct species were observed (30)). However, diversity in mechanical responses of GQs formed in longer telomeric DNA is yet to be explored.

In this study, we employed smFRET to probe GQ formation of human telomeric DNA, spanning five and six T_2_AG_3_ repeats. Using guanine to thymine (G to T) mutation at the central G of a T_2_AG_3_ repeat, which is known to restrict the participation of the repeat in GQ formation (24,30), we created variants without positional multiplicity to help interpret the data. Using smFRET and its combination with optical tweezers, which we refer to as fluorescence-force spectroscopy, we found evidence for preferential formation of GQs at the 3’ end of five-repeat telomeric DNA, with GQs formed at the 5’ end or at an internal position representing minority populations. Such GQs formed at the 3’ end unravel non-cooperatively via strand slippage, contrary to cooperative unfolding of canonical GQs formed from four telomeric repeats (30). Furthermore, using a superhelicase-based vectorial folding assay, we show that a GQ form preferentially at the 3’ end even when it is allowed to fold starting from the 5’ side. Although constituting a minor population, GQs formed at the 5’ end of a five-repeat telomeric DNA show extreme mechanical stability and cannot be perturbed by forces of ∼ 28 pN. GQs preferentially formed at the 3’ end of six telomeric repeats and manifested similar mechanical properties. Overall, we found that an unassociated T_2_AG_3_ repeat has a streamlining effect on GQ conformations in that one or two mechanically distinct conformations are observed at a single location instead of six or more observed for GQs without an unassociated repeat.

## Materials and Methods

### DNA constructs

All DNA oligonucleotides were purchased from Integrated DNA Technologies. The five and six telomeric repeat sequences referred to as hTel28 and hTel34 respectively are as follows: 5’-TGGCGACGGCAGCGAGGC**GGG(TTAGGG)_4_T**/Cy3/T_17_TCGGGAGCGGACGCACGG-3’ (hTel28) and 5’-TGGCGACGGCAGCGAGGC**GGG(TTAGGG)_5_T**/Cy3/T_17_TCGGGAGCGGACGCACGG-3’ (hTel34), where the telomeric repeat motif is bold-faced. For the repeat site selective mutation studies, five hTel28 mutants were designed such as G<u>T</u>G(TTAGGG)_4_T (hTel28.1), GGG(TTAG<u>T</u>G)(TTAGGG)_3_T (hTel28.2), GGG(TTAGGG)(TTAG<u>T</u>G)(TTAGGG)_2_T (hTel28.3), GGG(TTAGGG)_2_(TTAG<u>T</u>G)(TTAGGG)T (hTel28.4) and GGG(TTAGGG)_3_(TTAG<u>T</u>G)T (hTel28.5), where the underline denotes mutation form G to T. A truncated GQ strand hTel16, where GGG(TTAGGG)_3_T was replaced with GGG(TTAGGG)_2_T, was also used in our experiments. For further probing GQ formation in hTel34, an internal position was labeled with Cy3 and referred to as hTel34-4: 5’-TGGCGACGGCAGCGAGGC**GGG(TTAGGG)_3_T**/Cy3/**(TTAGGG)_2_T**T_17_TCGGGAGCGGAC GCACGG-3’. A complementary stem strand of sequence 3’/Biotin/ACCGCTGCCGTCGCTCCG/Cy5/5’ was used to immobilize the telomeric sequences of interest onto the slide surface. A second complementary strand (λ-bridge) of sequence 3’ AGCCCTCGCCTGCGTGCCTCCAGCGGCGGG 5’ was used to bridge the hTel strand with the 12 nt COS site of the λ-DNA. The other end of the λ-DNA could be further annealed with a digoxigenin labeled strand of sequence 3’AGGTCGCCGCCC/dig/5’. The hTel strands were first annealed with the complementary stem in 1.1:1 ratio in a buffer containing 50 mM NaCl and 10 mM Tris-HCl, pH 8, at 95 ^o^C for 5 min, followed by slow cooling to room temperature. The final construct for single molecule experiments by total internal reflection microscopy (TIRF) was generated by adding the λ-bridge DNA to the above mixture in the ratio of 1.5:1 to the complementary stem strand followed by incubation with rotation at room temperature for an hour.

For integrated smFRET-optical tweezers assay, λ-DNA (16 nM, New England Biolabs) was first heated in presence of 120 mM Na^+^ at 80 ^o^C for 10 min and then quenched on ice for 5 min. The hTel constructs and BSA were added to the λ-DNA at a final concentration of 8 nM and 0.1 mg/mL respectively and incubated/rotated at room temperature for 2-3 h. The dig-strand was next added to a final concentration of 200 nM and then incubated with rotation at room temperature for 1 h.

### Sample assembly

In order to eliminate non-specific surface binding, coverslips and quartz slides/glass slides used in our experiments were passivated with polyethylene-glycol (PEG) (a mixture of mPEG-SVA and biotin-PEG-SVA, Laysan Bio) (33). The PEGylated surfaces of the slides and coverslips were further sandwiched to form imaging chambers. For TIRF experiments, ∼ 30 pM hTel construct(s) were immobilized on the PEGylated surface via biotin-neutravidin interaction and imaged in a buffer containing 20 mM Tris-HCl pH 8, 0.8 % w/v D-Glucose [Sigma], 165 U/mL glucose oxidase [Sigma], 2170 U/mL catalase [Roche], 3 mM Trolox [Sigma] and pre-determined amount of NaCl or KCl.

For integrated smFRET-optical tweezers experiments, a similar imaging chamber was incubated in blocking buffer containing 10 mM Tris HCl pH 8, 50 mM NaCl, 1 mg/mL BSA [NEB] and 1 mg/mL tRNA [Ambion] for 1 h. The DNA constructs were immobilized on the surface at ∼ 10 pM concentration, via biotin-neutravidin interaction. Next, anti-digoxigenin coated polystyrene beads (Polysciences) were attached to the immobilized construct by incubation in a buffer containing 10 mM Tris HCl pH 8 and 50 mM NaCl for 30 min. Finally, data were acquired in an imaging buffer consisting of 50 mM Tris-HCl pH 8, 0.8 % w/v D-Glucose [Sigma], 0.5 mg/mL BSA [NEB], 165 U/mL glucose oxidase [Sigma], 2170 U/mL catalase [Roche], 3 mM Trolox [Sigma] and pre-determined amount of NaCl/KCl.

### Fluorescence-force spectroscopy

An integrated single molecule fluorescence-optical trap instrument was recently developed in our lab to probe conformational changes of biomolecular systems under tension (34,35). In short, an optical trap was formed by an infrared laser (1064 nm, 800 mW (maximum average power), EXLSR-1064-800-CDRH, Spectra-Physics) through the back port of the microscope (Olympus) on the sample plane with a 100X immersion objective (Olympus). Tension was applied on the sample tethers by translating the piezo-electric stage that holds the microscope and the applied force was read out via position detection of the tethered beads using a quadrant photodiode (UDT/SPOT/9DMI). The instrument was calibrated as described previously (34,35). A confocal excitation laser (532 nm, 30 mW (maximum average power), World StarTech) was used to scan the sample via a piezo-controlled steering mirror (S-334K.2SL, Physik Instrument). Two avalanche photodiodes were used to record the fluorescence emission, filtered from infrared laser by a band pass filter (HQ580/60 m, Chroma) and excitation by a dichroic mirror (HQ680/60 m, Chroma.

### Data acquisition

In the absence of force, a prism-type total internal reflection microscope, with 532 nm laser excitation and back-illuminated electron-multiplying charge-coupled device camera (iXON, Andor Technology, South Windsor, CT) was used to detect smFRET signals (33). The donor and acceptor intensities, i.e. *I*_D_ and *I*_A_ respectively were corrected for background signals and crosstalk and used to estimate smFRET efficiency *E* using *I*_A_/(*I*_A_+*I*_D_). *E* histograms were constructed by averaging the first ten data points of each molecule’s time trace.

A detailed data acquisition procedure for integrated single molecule force-fluorescence spectroscopy has been described by Hohng *et al* (34). Briefly, a tethered bead was trapped and its origin was determined by stretching the tether along opposite directions along x and y-axes. The trapped bead was next moved from its origin by 14 μm and fluorescence spot on the tether located by scanning with the confocal. All unfolding and/or subsequent refolding experiments were performed by translating the microscope stage at a speed of 455 nm/s between 14 μm and 16.8-17.2 μm. Fluorescence emission from the tethered molecule was detected simultaneously with the stage movement, 20 ms after each step in the stage movement. Similar to TIRF-based smFRET experiments, force-fluorescence data was obtained in the imaging buffer containing 50 mM Tris-HCl pH 8, 0.8 % w/v D-Glucose [Sigma], 0.5 mg/mL BSA [NEB], 165 U/mL glucose oxidase [Sigma], 217 U/mL catalase [Roche], 3 mM Trolox [Sigma] and pre-determined amount of NaCl/KCl.

The *f_unfold_* histograms were analyzed and the free energy parameters such as the transition distances to unfolding etc were predicted with reference to the Dudko-Szabo model (ν=1/2). These derived parameters (*Δx*^‡^, *τ_u_*(0), and *ΔG*^‡^) were then used to reconstruct the force profiles (36,37).

### Circular Dichroism (CD) Spectroscopy

CD spectra of the human telomeric oligonucleotides were recorded on an Aviv-420 spectropolarimeter (Lakewood, NJ, USA), using a quartz cell of 1 mm optical path length. The oligonucleotides were diluted to 20 μM in a buffer containing 20 mM Tris pH 8 and appropriate concentration of K^+^/Na^+^ ions. Before each measurement, the oligonucleotides were heated to 90 ^o^C for ∼ 5 min and slowly cooled to room temperature, to avoid formation of intermolecular structures. An average of three scans was recorded between 220 and 320 nm at room temperature. The spectra were corrected for baseline and signal contributions from the buffer.

### Vectorial GQ folding using a superhelicase Rep-X

In order to eliminate end-effects of the sequence on Rep-X unwinding and GQ folding, we introduced additional 10 nt at the 3’ end of hTel28: 5’-TGGCGACGGCAGCGAGGC**GGG(TTAGGG)_4_T**/Cy3/GGACGCACGG-3’. The hTel28 was then annealed to its complementary sequence in the ratio of 1:1.1 at 95 ^o^C for 5 mins, in a buffer containing 50 mM NaCl and 10 mM Tris-HCl, pH 8, followed by slow cooling to room temperature. The duplexes were then immobilized on the PEG-passivated surface. 50 nM Rep-X was then incubated in a loading buffer (10 mM Tris-HCl, pH 8, 10 % glycerol, 1 % BSA) for 2 min. The unbound Rep-X was washed off simultaneously and the unwinding reaction was initiated by adding the unwinding buffer (10 mM Tris-HCl, pH 8, 100 mM KCl, 2.5 mM MgCl_2_, 1 mM ATP [ThermoFisher Scientific], 10 % glycerol, 1 % BSA). For characterization of the maiden folded GQ conformation, the unwinding reaction was quenched after a minute and images were acquired for smFRET analysis. For the real-time observations, imaging was started a few seconds before addition of the unwinding buffer. A detailed protocol has been described by Hua *et al* (38).

## Results

### SmFRET analysis of five repeats human telomeric DNA

Our hTel28 construct consists of a central 28 nt long human telomeric DNA sequence spanning five hexanucleotide repeats, (GGG(TTAGGG)_4_T), flanked by two duplex handles at the 5’ and 3’ ends. CD spectrum of hTel28 in 100 mM K^+^ showed a positive peak at ∼ 289 nm with shoulders at 265 and 255 nm and a negative peak at 235 nm, suggestive of hybrid GQs (Figs. 1a, S1a) (39). The construct was immobilized on to the PEG-passivated surface via the duplex stem at the 5’ end whereas a 3’end partial duplex served as a tether to the λ-DNA for use in fluorescence-force spectroscopy. The donor (Cy3) and acceptor (Cy5) fluorophores were placed adjacent to the ends of the telomeric sequence such that FRET efficiency, *E*, reflects the end-to-end distance of the telomeric sequence (Fig. 1b).

**Fig. 1:**
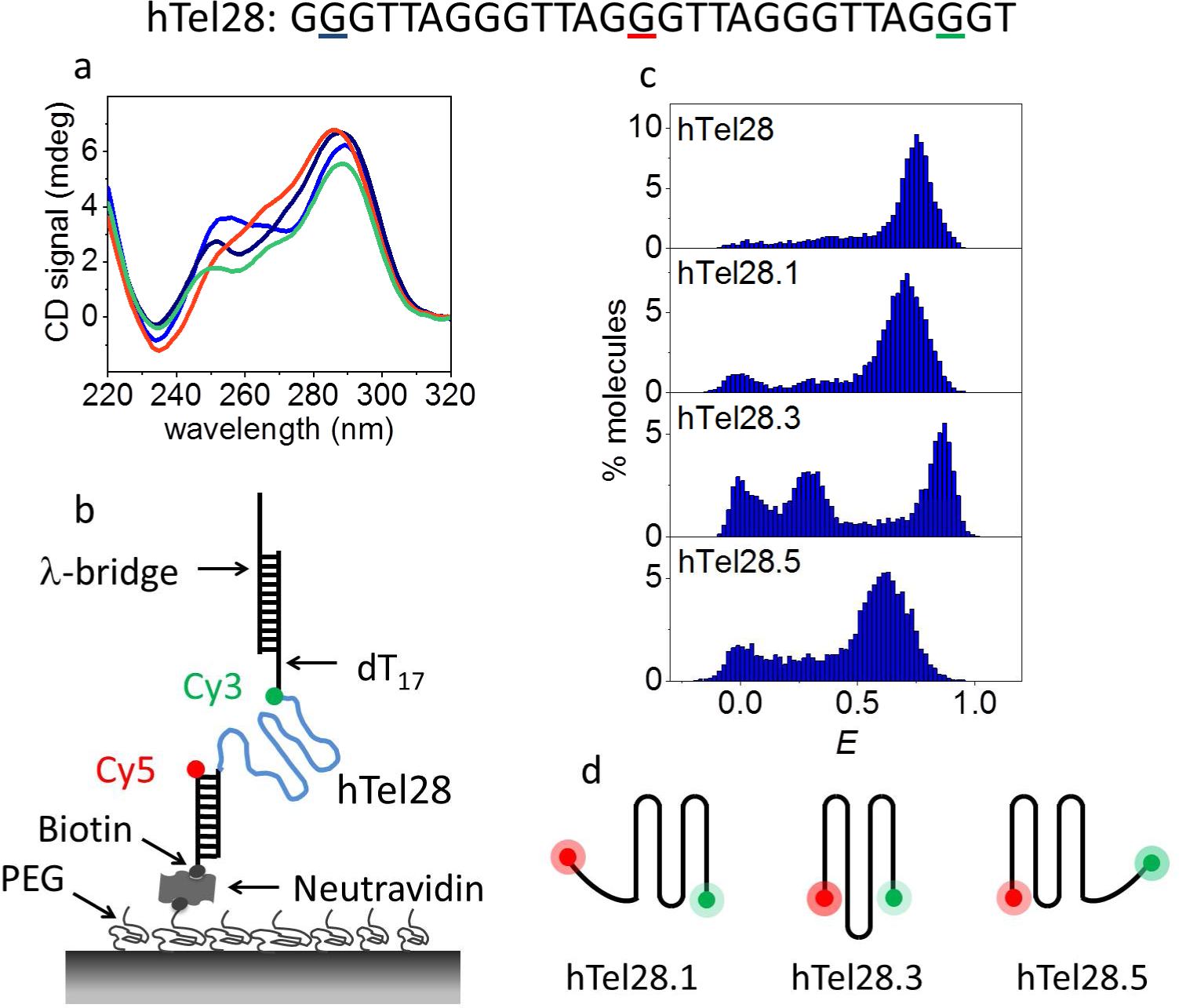
Conformational analysis of a five-repeat telomeric DNA (hTel28). (a) CD spectra of hTel28 (blue), hTel28.1 (navy blue), hTel28.3 (red) and hTel28.5 (green) in 100 mM K^+^. The G to T mutation sites in hTel28 corresponding to hTel28.1, hTel28.3 and hTel28.5 are underlined on the sequence with navy blue, red and green respectively. (b) Schematic diagram of hTel28 construct for smFRET. The 5’ extension to the 28 nt long human telomeric repeat GGG(TTAGGG)_4_T was annealed to a 18 nt long biotinylated strand and immobilized on a PEG-passivated quartz surface through biotin-neutravidin interaction. The 5’ end of the biotinylated strand is labeled with Cy5 (acceptor). FRET was measured between Cy3 and Cy5. The main hTel28 strand is labeled with Cy3 (donor) at the 3’ end of the telomeric repeat and is followed by dT_17_ and an 18 nt extension that is annealed to 30 nt long λ-bridge. (c) *E* histograms of hTel28, hTel28.1, hTel28.3 and hTel28.5 in 100 mM K^+^ concentration. (d) Schematic representation of GQ formation after site-specific G to T mutation of hTel28.

Formation of secondary structures was first examined through smFRET as a function of K^+^ concentration (Fig. S1b). The peak centered at *E* = 0 represents molecules with a missing or inactive acceptor, and hence can be ignored. In the absence of K^+^, we observed a peak centered at *E* ∼ 0.16. Transition into secondary structures commenced between 2 and 10 mM K^+^, as suggested by emergence of broad peaks at *E* ∼ 0.3 and ∼ 0.69 at 10 mM K^+^, which then resulted in a major folded population at *E* ∼ 0.75 at 100 mM K^+^ (Figs. 1c, S1b). The populations of low and high *E* values were attributed to unfolded and folded conformations, respectively. Single molecule trajectories showed no *E* transitions during our observation time up to ∼ 90 s at a time resolution of 30 ms (Figs. S1d).

In a five-repeat telomeric sequence, GQs can potentially form at the 3’ or 5’ ends by association of four consecutive T_2_AG_3_ repeats (3’ and 5’ GQs respectively), or with an inner long loop bearing the second, third or fourth T_2_AG_3_ segment (long-loop GQs). Therefore, at least five different GQ structures can be conceived depending on which repeat is excluded (Fig. S2). In order to shed light on the conformational landscape, mutations can be introduced in biomolecules to selectively depopulate certain species from a heterogeneous population (24,29,40). Previous studies have also shown that guanine to thymine substitution of the central G of a hexanucleotide repeat T_2_AG_3_ precludes its participation in GQ formation in 100 mM K^+^ or Na^+^ (24,30) guanine of one T_2_AG_3_ segment in hTel28 by thymine to selectively reduce the possibility of that segment becoming part of a GQ. Five such variants of hTel28 were designated as hTel28.1, hTel28.2, hTel28.3, hTel28.4 and hTel28.5 based on the position of the segment bearing G to T mutation, counting from the 5’ end. For example, hTel28.1 would fold into a 3’ GQ because the mutated 5’ most repeat would not participate in a GQ under physiological ionic conditions from the 5’ end. For example, hTel28.1 would fold into a 3’ GQ because the mutated 5’ most repeat would not participate in a GQ under physiological ionic conditions.

CD spectra of the hTel28 variants in 100 mM K^+^ showed signatures of hybrid GQs as was the case for hTel28 itself (Fig. 1a) (39). However, they cannot individually fully recapitulate that of hTel28, suggesting co-existence of multiple species in hTel28. Folding of the hTel28 variants into secondary structures was modulated by cationic concentrations (data not shown) and in 100 mM K^+^, a major high *E* population with peaks centered at ∼ 0.71, 0.85, 0.85, 0.86 and 0.62 was observed for hTel28.1, hTel28.2, hTel28.3, hTel28.4 and hTel28.5, respectively (Fig. S1c). Previous smFRET studies with a 22 nt GQ-forming telomeric sequence (hTel22) harboring four repeats reported a major *E* peak centered at ∼ 0.88 in 100 mM K^+^ (30,32), similar to the peaks at *E* ∼ 0.85 observed for hTel28.2, hTel28.3 and hTel28.4. Therefore, we tentatively assign *E* ∼ 0.85 subpopulation of hTel28 to long-loop GQs where a telomeric repeat at the 2^nd^, 3^rd^ or 4th position becomes a part of a loop (Fig. S2). On the other hand, GQs formed from four consecutive repeats mimicked by hTel28.1 and hTel28.5 would leave a 6 nt single stranded portion, adding to the separation between the fluorophores (Fig. S1c), consistent with the lower *E* values observed for hTel28.1 and hTel28.5 compared to the long-loop GQs. The hTel28 variants were very stable as evidenced by steady *E* values (Fig. S1d).

Among the variants, hTel28.1 appears closest to hTel28 in the major smFRET population’s peak position, suggesting a preferential formation of GQ at the 3’ end of a five-repeat telomeric sequence (Figs. 1c and S1c). However, given the significant overlap with the *E* distributions of the other four hTel28 variants and hTel28, 5’ end GQs and long-loop GQ may exist as minor populations (Figs. 1c and S1d). In fact, using the peak centers and widths from *E* histograms of hTel28 variants, we were able to resolve three non-interconvertible subpopulations in hTel28 with *E* values of ∼ 0.66, 0.75 and 0.85 accounting for ∼18 % (37 out of 200 events), 64 % (128 events) and 18 % (35 events) of the recorded events, respectively (Fig. S1c).

### Vectorial folding of hTel28, mimicking telomerase action

Our smFRET studies under refolding conditions suggest predominant GQ formation at the 3’ end of hTel28s. However, because a telomerase adds one repeat at a time to the 3’ end of a telomeric DNA under a physiological situation of telomere elongation, a GQ may first form on the 5’ side during telomere elongation. To address this point, we employed a highly processive superhelicase Rep-X to mimic the vectorial nature of telomerase action (38,41,42). In a previous study, we used this 5’ to 3’ DNA helicase to unwind RNA/DNA heteroduplex and to reveal the RNA strand in the direction of transcription (5’ to 3’) at the speed of transcription to mimic co-transcriptional RNA folding (38). Here, we annealed hTel28 to its complementary C-rich DNA and Rep-X was loaded through the 3’ overhang in the absence of MgCl_2_ and ATP (Fig. 2a). Duplex unwinding was then initiated by addition of MgCl_2_ and ATP. The helicase translocates in the 3’ to 5’ direction along the C-rich DNA strand, unwinding the duplex and releasing the G-rich DNA in the 5’ to 3’ direction, which is the direction of telomere extension. The disengaged G-rich hTel28 can fold into GQ as it is being revealed starting from the 5’ side (Fig. 2a). GQ folding in nascent telomeric DNA synthesized during telomerase extension has been directly observed by Jansson *et al* (43).

**Fig. 2:**
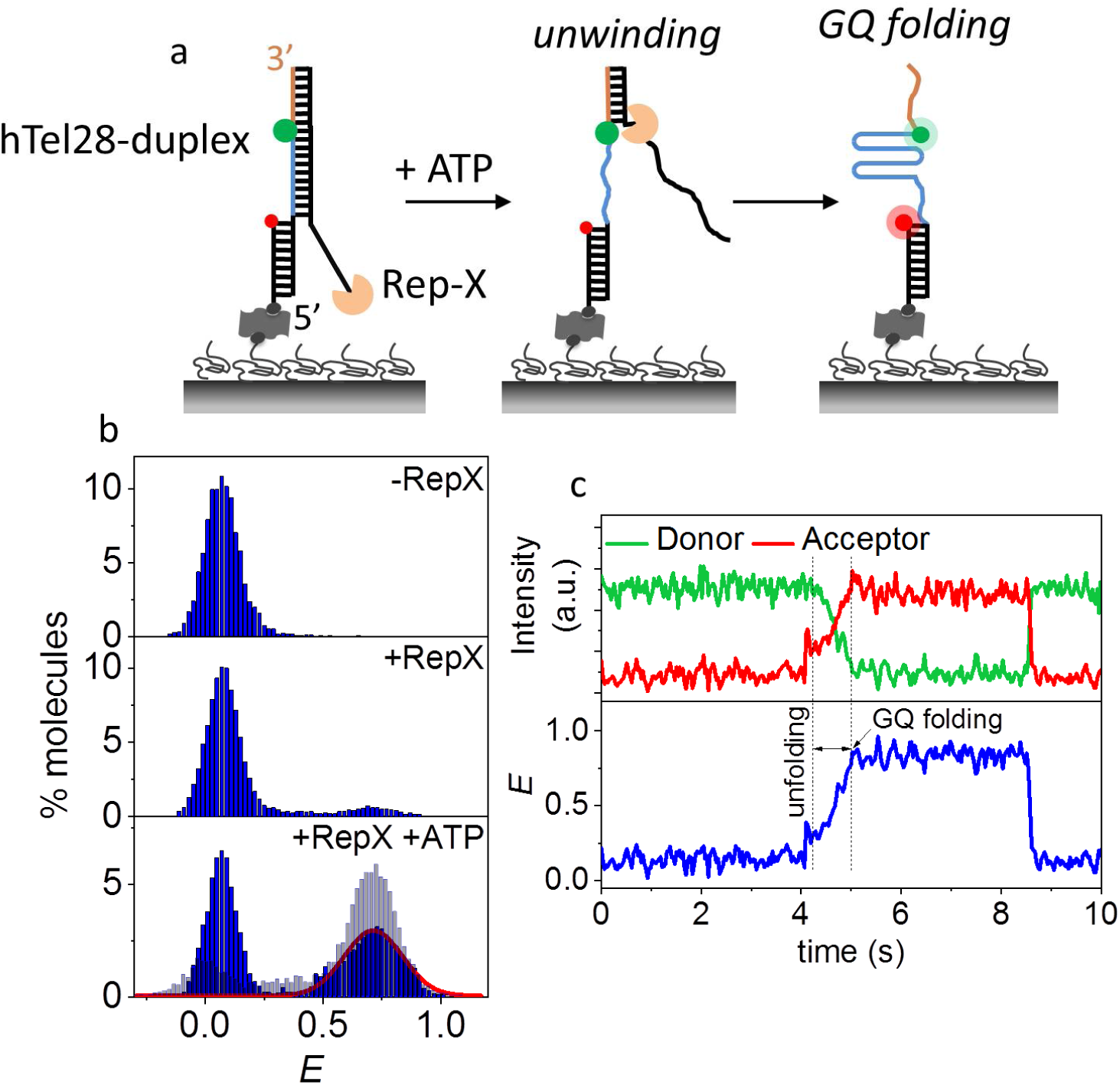
Vectorial folding of hTel28. (a) Schematic of unfolding of a hTel28-duplex via Rep-X and subsequent folding of hTel28 into GQ. (b) *E* histograms of hTel28-duplex in the Rep-X unbound (top) and bound (middle) states, and after Rep-X unwinding of the duplex in presence of ATP and MgCl_2_ (bottom). The red curve represents Gaussian fit to the folded GQ population. *E* histogram of hTel28.1 in the unwinding buffer is superimposed in light blue (bottom). (c) Representative single molecule trajectory showing unwinding of the duplex and folding of GQ (30 ms integration time).

In the fully annealed state, hTel28-duplex showed a FRET peak at *E* ∼ 0.07 (Fig. 2b). Rep-X mediated unwinding reaction was quenched after a minute by washing away ATP and the resulting smFRET distribution showed that ∼ 47 % of molecules fold into a GQ with a major peak at *E* ∼ 0.73 (Fig. 2b). The rest stayed at *E* ∼ 0.07 probably because these molecules did not get unwound. A control experiment performed with a slowly hydrolysable ATP analog, AMP-PNP, showed only ∼ 5 % folded population (Fig. S3d). Under identical unwinding buffer conditions, major peaks centered at *E* ∼ 0.72 and 0.62 were observed in hTel28.1 and hTel28.5, respectively (Fig. S5a). Hence, the major state observed at *E* ∼ 0.73 was attributed to 3’ GQs.

Real-time single molecule trajectories during unwinding typically showed gradual increases in *E* to ∼ 0.73 over ∼ 0.7 s without pausing at intermediate FRET states (Figs. 2c, S3b and c). Such behavior was observed in ∼ 64 % (74 of 116 events) of the folded GQs. This suggests that GQ forms at the 3’ end even during vectorial folding. 5’ GQ may form transiently but if so, it is not long-lived enough to be detected as a clear intermediate. About 22 % (26 events) of the folded GQs showed *E* ∼ 0.62, suggestive of 5’ GQs as a minority population. Thus, although vectorial unwinding of hTel28-duplex via Rep-X exposes the T_2_AG_3_ segments successively from the 5’ end, GQs are formed preferentially at the 3’ end.

### Dynamics of hTel28 under tension

For fluorescence-force spectroscopy, the hTel28 construct was annealed with λ-DNA via the λ-bridge prior to its immobilization on the PEG-passivated surface. The other end of the λ-DNA was tethered to an optical trapped bead (Fig. 3a). The molecule under investigation was stretched and relaxed by translating the sample stage at a speed of 455 nm/s. In a typical experiment, the applied force was increased from ∼ 0.3 pN to ∼ 28 pN in ∼ 6.5 s, followed by relaxation at the same speed. The donor and acceptor fluorescence intensities were recorded as a function of force in each pulling cycle and this was repeated until fluorophore photobleaching or tether rupture. Data were collected from at least 20 different molecules under each condition.

**Fig. 3:**
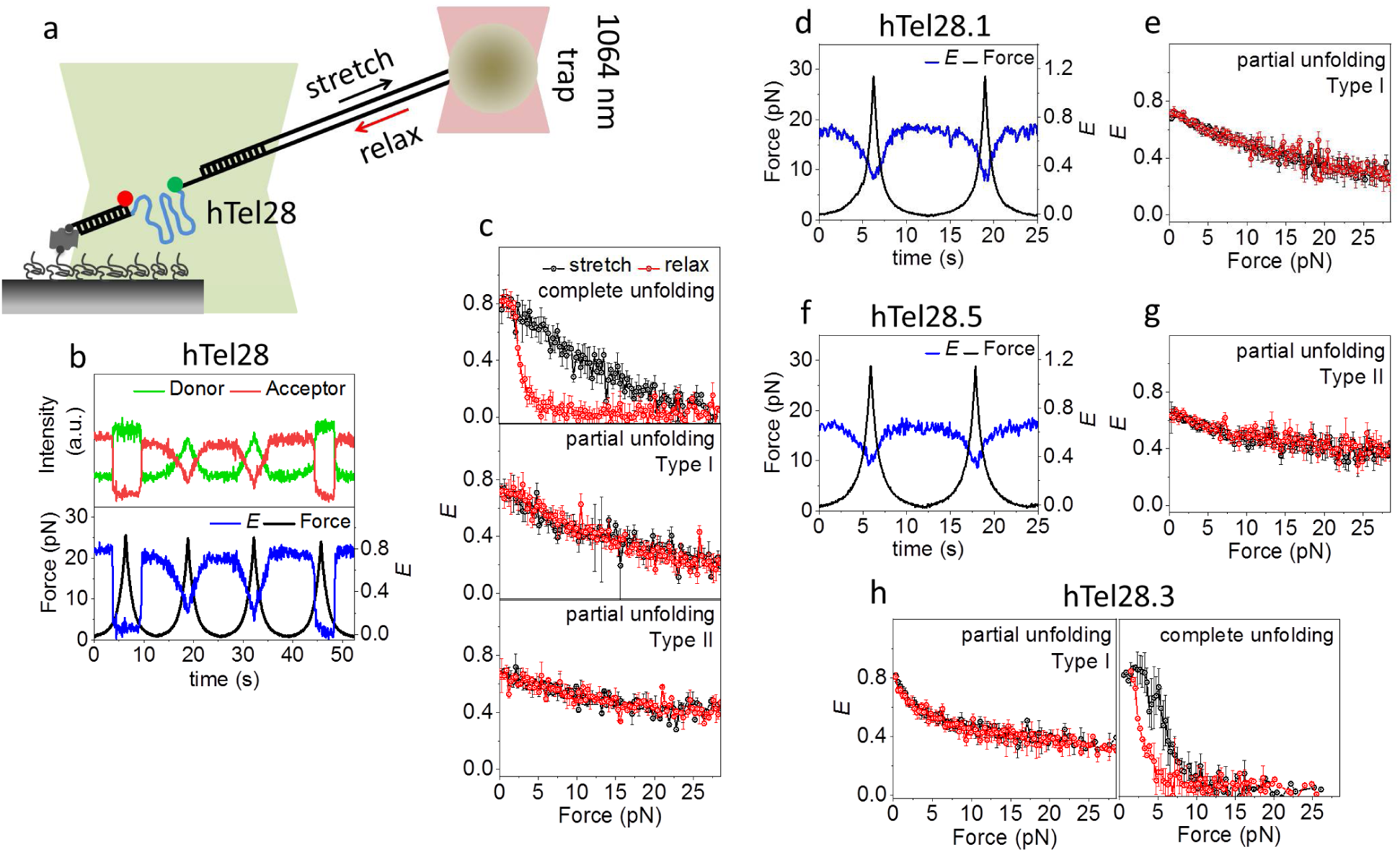
Conformational dynamics of five human telomeric repeats under tension in 100 mM K^+^. (a) Schematic of integrated fluorescence-force spectroscopy assay: The hTel28 strand is annealed to a biotinylated strand and immobilized on a neutravidin coated quartz surface. The other end is connected to an optically trapped 1 μm diameter bead through a λ-DNA. Force was applied by translating the microscope stage at a speed of 455 nm/s. FRET was measured between Cy3 (donor) and Cy5 (acceptor) as a function of force. (b) A representative single molecule time trace of donor and acceptor intensities and corresponding *E* of hTel28. (c) Average *E* vs force response from molecules undergoing complete (*N*=58, top) partial (I) (*N*=150, middle), and partial (II) (*N*=55, bottom) unfolding under tension. (d and f) Representative *E* time traces of hTel28.1 (d) and hTel28.5 (f) during two pulling cycles. (e, g and h) Average *E* vs force response of hTel28.1 (e), hTel28.5 (g) and hTel28.3 (h). All single molecule trajectories were collected at a time resolution of 20 ms. Error bars represent standard errors.

Diverse conformational changes of hTel28 in 100 mM K^+^ were observed under tension. The extent of hTel28 unraveling, as judged by the lowest *E* values achieved, varied across different stretching cycles of a single molecule (Fig. 3b) and could be broadly categorized as complete unfolding (22 %, 58 out of 263 events), Type I partial unfolding (∼57 %, 150 events) and Type II partial unfolding (∼21 %, 55 events). Typically, complete unfolding of a molecule was characterized by stable *E* ∼ 0.8 up to a certain force, followed by an abrupt decrease in *E* to ∼ 0.1 (Figs. 3b, cycles 1 and 4, and S4a). Further gradual decreases from *E* ∼ 0.1 suggests stretching of an already unfolded molecule. The subsequent relaxation event, similarly, showed an abrupt recovery to the initial *E* value. The unfolding (*f_unfold_*) and refolding forces (*f_refold_*) were defined as the force(s) corresponding to midpoint(s) of the unfolding and refolding transitions. Diverse *f_unfold_* values observed across different pulling cycles could be fit to two force clusters centered at ∼ 7.5 and 17 pN (Figs. S4b and c). Refolding, on the other hand, predominantly occurred at lower forces, between 1 and 7 pN (Fig. S4d).

The most common was Type I partial unfolding, defined by stable *E* values of ∼ 0.75 up to ∼ 2 pN, followed by a gradual decrease to ∼ 0.25 at ∼ 28 pN (Fig. 3b, cycles 2 and 3). On relaxation, the unraveling pathway was retraced without any significant hysteresis (Fig. 3c). Type II partial unfolding also showed reversible *E* changes but with a different range, from ∼ 0.68 to ∼ 0.37 (Fig. 2g).

Interestingly, a single molecule of hTel28 under tension can interchange between complete and Type I partial unfolding. For example, Fig. 3b shows complete and partial unfolding in consecutive pulling cycles 1 and 2 and vice versa in the cycles 3 and 4. Overall, switching from complete to Type I partial unfolding in consecutive cycles and vice versa were observed in ∼ 33% (19 out of 58 events) and ∼ 16 % (24 out of 150 events) pulling cycles, respectively. However, Type II partial unfolding events rarely switched to other types of unfolding in the subsequent cycle (4 events, ∼ 7 %).

### Conformational dynamics of hTel28 variants under tension

We next examined the hTel28 variants under tension in order to gain mechanistic insight into GQ formation in five-repeat telomeric DNA. In 100 mM K^+^, hTel28.1 showed a stable *E* of ∼ 0.72 up to applied forces of ∼ 2 pN, followed by a gradual decrease in *E* to ∼ 0.25 at a force of ∼ 28 pN (Figs. 3d, e and S5c). It retraced the same *E* vs force during relaxation (Fig. 3e) and subsequent pulling cycles yielded similar stretching-relaxation behavior. hTel28.5 also showed reversible changes in *E* upon stretching and relaxation. However, the *E* values changed between 0.35 and 0.62, which is narrower in range than hTel28.1 which ranged between 0.25 and 0.72 (Figs. 3f, g and S5c). Conformational dynamics of hTel28.1 and hTel28.5 under tension therefore are very similar to Type I and Type II partial unfolding observed in hTel28, respectively. Therefore, we propose that 3’ GQs, mimicked by hTel28.1, show Type I partial unfolding and 5’ GQs, mimicked by hTel28.5, show Type II partial unfolding.

We next studied the conformational dynamics of hTel28.3 as a representative of long-loop GQs. Because the hTel28.3 harbors a G to T mutation in the innermost T_2_AG_3_ segment, a GQ would form preferentially with the four outer repeats with an inner 9 nt long loop (Fig. S5a). Complete and abrupt unfolding characterized by *E* drop from ∼ 0.83 to ∼ 0.15 and subsequent single-step refolding at lower forces accounted for ∼ 48 % (50 out of 104 events) of the pulling cycles (Figs. 3h and S5d-g). The *f_unfold_* values clustered around an average value of ∼ 6 pN, close to the ∼ 7.5 pN *f_unfold_* cluster observed in complete unfolding of hTel28 (Figs. S5b and d). The remaining 52 % (54 events) showed gradual, reversible changes in *E* from ∼ 0.83 to ∼ 0.4, similar to Type I partial unfolding of hTel28 (Fig. 3h).

In all, mechanical behavior of hTel28 was recapitulated by the hTel28 variants studied. We can attribute complete unfolding to long-loop GQs, and attribute Type II partial unfolding to 5’ GQs. Type I partial unfolding was observed in both long-loop GQs and 3’ GQs. Interestingly, among the variants, mechanical heterogeneity could be noted only in long-loop GQs (hTel28.3). Combining these results with zero-force smFRET studies, we propose that in 100 mM K^+^, the 3’ GQs constitute the preferred conformation of hTel28.

### Effect of an unassociated G-rich segment at the 3’ end of GQ

We previously classified diverse mechanical behavior of hTel22 that contain four telomeric repeats into (1) abrupt and complete unfolding, (2) partial unfolding and (3) an ultrastable population that could not be perturbed by forces up to ∼ 28 pN (30). GQs formed at the 3’ or 5’ ends of hTel28 would have a 6 nt overhang to an hTel22-like GQ core. Therefore, an ultrastable GQ with a pendant 6 nt tail would show gradual *E* decreases as the ssDNA tail is stretched. Thus, the reversible *E* changes between ∼ 0.68 and ∼ 0.39 under applied forces shown by 5’ GQs and classified as Type II partial unfolding of hTel28 may be due to an ultrastable GQ population. On the other hand, Type I partial unfolding which entails a greater degree of unraveling (Fig. S5c) can be due to partial unfolding via mutual slippage of the G-rich strands as proposed to occur in hTel22 (30). Interestingly, Type I partial unfolding, although dominant in hTel28, was observed only in ∼ 11 % of hTel22 molecules (30). Hence, in order to investigate the effect of a short addendum at the 5’ end of GQ, we substituted the 6 nt T_2_AG_3_ segment at the 5’ end with (dT)_6_ (hTel28.1T). Thus, hTel28.1T serves as a 22 nt long GQ forming sequence with a pendant (dT)_6_ at its 5’ end.

hTel28.1T folded in a K^+^ concentration-dependent manner (Fig. S6a), showing a major population at *E* ∼ 0.77 (Figs. 4a and S6a). The *E* values are similar between hTel28.1T, hTel28 and hTel28.1, further suggesting that a GQ forms predominantly at the 3’ end of a five-repeat telomeric sequence. Under tension in 100 mM K^+^, hTel28.1T showed three qualitatively different types of behavior. For example, a molecule can gradually stretch to *E* ∼ 0.47 from an initial *E* ∼ 0.77 and back, likely as a result of stretching of the 6 nt ssDNA next to an ultrastable GQ (Figs. 4b (cycle 1) and S6c) (44). The range of *E* values is comparable with that observed in Type II partial unfolding, supporting our assignment of hTel28.5 to an ultrastable GQ. In the second and third pulling cycles of Fig. 4b, we observed a gradual decrease in *E* from ∼ 0.77 due to ssDNA stretching, followed by abrupt unfolding (Fig. 4b). Refolding also occurred abruptly (Figs. 4b and d). This second type of behavior showed a wide range of unfolding forces (∼ 1 and 28 pN) (Figs. 4c and S6b), similar to cooperative unfolding observed for hTel22 (30). The *f_unfold_* values (131 events) can be grouped into three force clusters centered at ∼ 3.5, 10 and 20 pN (Fig. 4c). We also observed a third type of unraveling behavior, characterized by gradual and reversible changes from *E* ∼ 0.77 to ∼ 0.25 (24 events, ∼ 10 %) and reminiscent of Type I partial unfolding of hTel28 (Fig. S6d) (30). In all, we could recapitulate conformational dynamics of hTel22 with hTel28.1T. The stark contrast in mechanical behavior of hTel28.1 and hTel28.1T suggests that GQ conformations are affected by the unassociated G-rich strand at the 5’ end in a way that a (dT)_6_ cannot mimic.

**Fig. 4.**
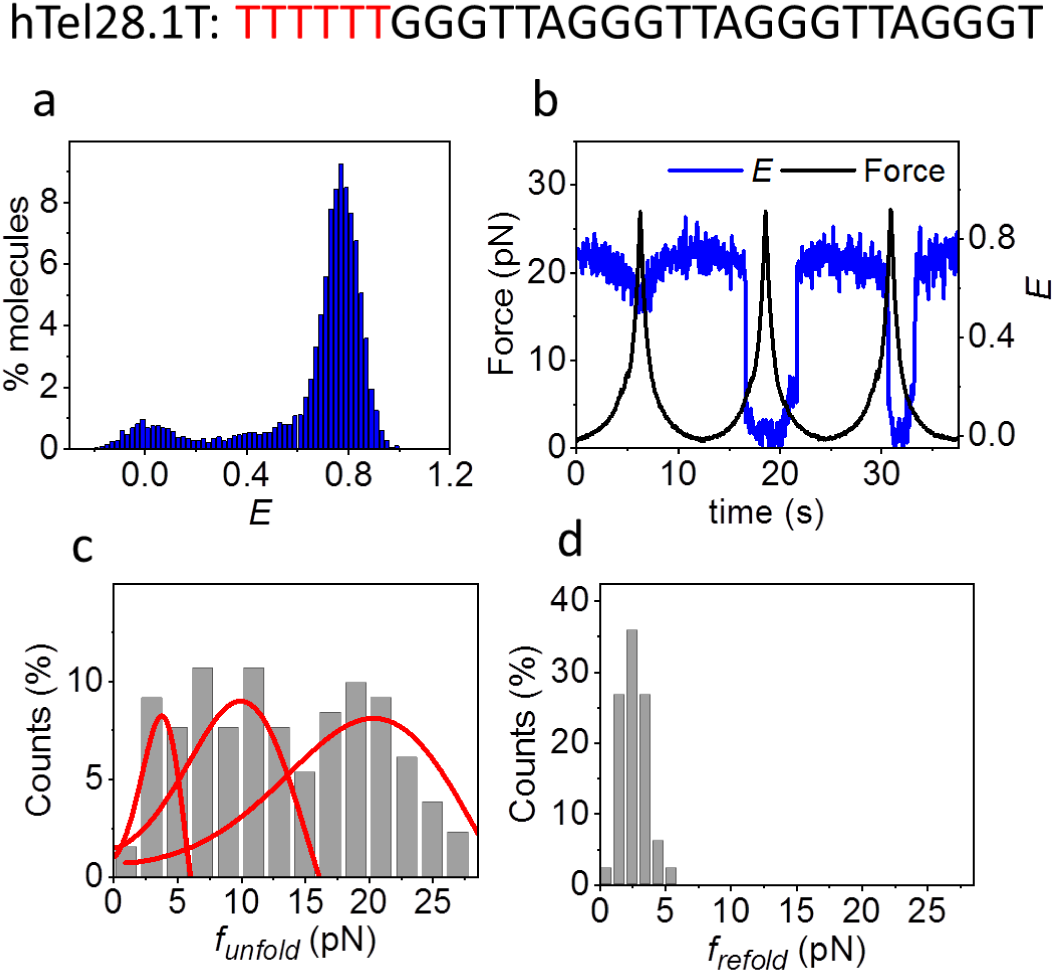
(a) *E* histogram of hTel28.1T in 100 mM K^+^, in the absence of force. (b) A representative *E* time trace of hTel28.1T during three pulling cycles. (c and d) Distributions of *f_unfold_* (c) and *f_refold_* (d). Only those cycles showing complete unfolding were included (*N*=131). The red curves represent the rupture force distributions predicted using the Dudko-Szabo model (36,37). Error bars represent standard errors.

### Formation of GQs in six-repeat human telomeric DNA

We next investigated the formation of GQs in six-repeat human telomeric DNA, hTel34 (Fig. 5a). Like hTel28, CD spectra of hTel34 in 100 mM K^+^ shows signatures of hybrid GQs (Fig. 5b) (39). Secondary structure formation commenced at < 10 mM K^+^ (Fig. S7a), and in 100 mM K^+^, we observed two non-interconvertible (within our observation time) major populations, centered at *E* ∼ 0.54 and 0.71, that we designate as *E*_1_ and *E*_2_ states (Fig. 5c, Fig. S7b). A minor population at *E* ∼ 0.33 (*E_0_*) is similar in mechanical response to (dT)_34_, an unstructured ssDNA of the same length (Fig. 5i and 5j), and therefore is likely an unfolded population.

**Fig. 5:**
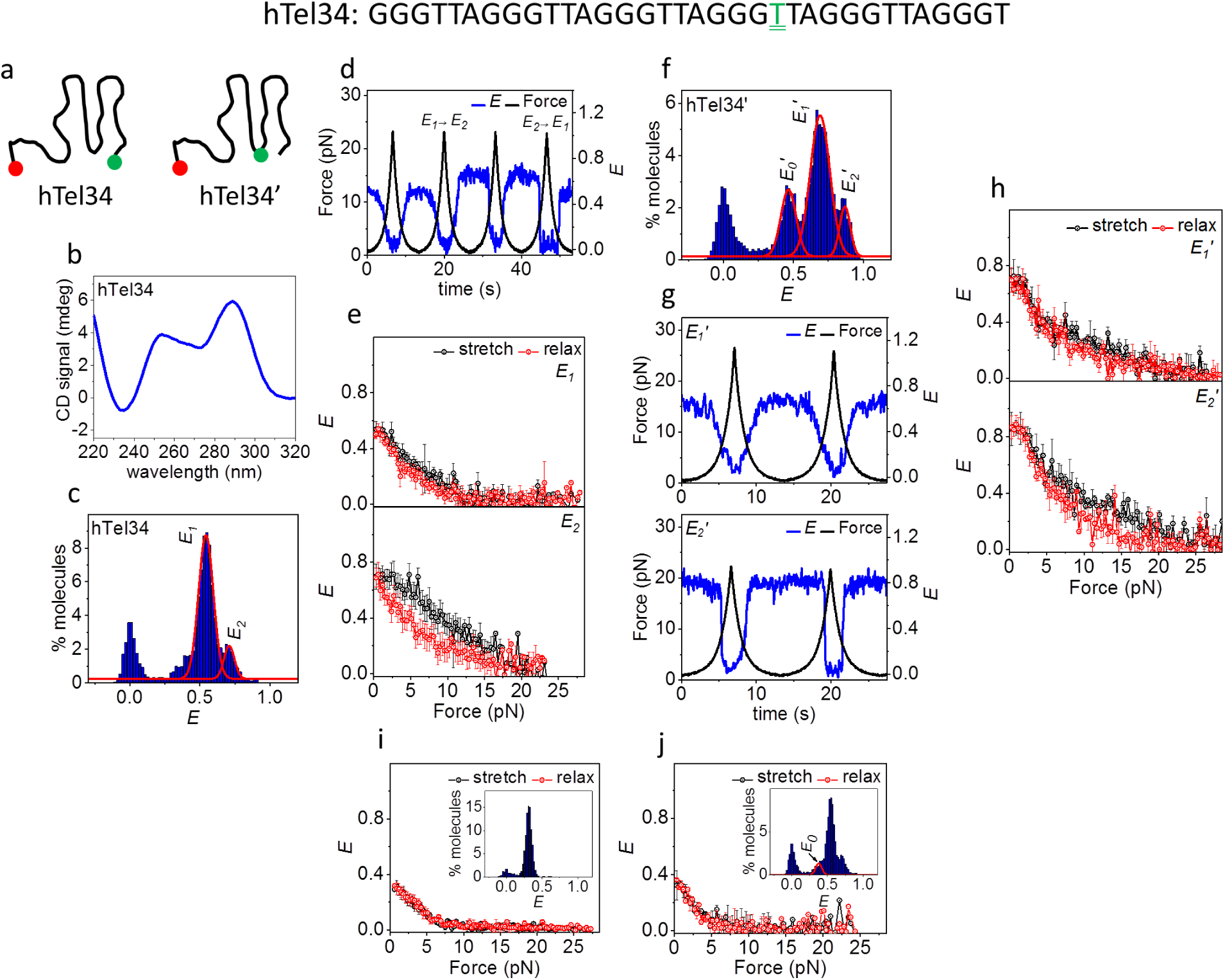
Conformational analysis of hTel34 in 100 mM K^+^. (a) Schematic representations of hTel34 and htel34’. T_22_ bearing the donor dye in hTel34’ is doubly underlined in green. (b) CD spectra of hTel34 in 100 mM K^+^. (c) *E* histogram of hTel34 in the absence of force. The red curves denote *E_1_* and *E_2_* populations. (d) A representative *E* time trace of hTel34 during four pulling cycles. (e) Average *E* vs force plots of *E_1_* (*N*=89, top) and *E_2_* (*N*=48, bottom). (f) *E* histogram of hTel34’ in the absence of force. The red curves denote the *E_0_’*, *E_1_’* and *E_2_’* populations. (g) Representative *E* time traces of *E_1_’* (top) and *E_2_’* (bottom) of hTel34’ during two pulling cycles. (h) Average *E* vs force plots of *E_1_’* (*N*=128, top) and *E_2_’* (*N*=58, bottom). (i and j) Average *E* vs force response of (dT_34_) (i) and *E_0_* of hTel34 (j). The corresponding *E* histograms in the absence of force are shown in inset. Error bars represent standard errors.

Under tension, *E*_1_, which is the dominant population, unraveled gradually and reversibly (89 out of 137 events, ∼ 65 %, Figs. 4d and e) whereas *E_2_* unfolding occurred abruptly (48 events, Fig. 5d). *f_unfold_* for *E*_2_ showed a wide distribution (1 - 20 pN) that could be fitted to two clusters centered at ∼ 7.5 and 16 pN (Figs. S7c and d) and refolding occurred at lower forces (Figs. 5e and S7d). Switch from *E_2_* to *E_1_* after a pulling cycle accounted for ∼ 29 % (14 out of 48 events, Fig. 5d (cycle 4)) of the events originating at *E_2_* whereas a *E_1_* to *E_2_* switch was observed in ∼ 13 % (12 out of 89 events, Fig. 5d (cycle 2)) of the events initiating from *E_1_*.

With two additional T_2_AG_3_ repeats, hTel34 can form 15 GQ structures that differ in which four of the six repeats are chosen (Fig. S2). In order to gain information on its internal architecture we placed the donor fluorophore at the thymine in the 22^nd^ position (hTel34’, Fig. 5a) so that FRET reports on the conformation of the first four repeats, counting from the 5’ end. We have previously shown that fluorescent labels do not significantly alter the folded states of GQs (30). At 100 mM K^+^, the smFRET distribution showed three major non-interconvertible (within our observation time) populations centered at *E* ∼ 0.47 (*E_0_’*), 0.68 (*E_1_’*) and 0.86 (*E_2_’*) (Figs. 5e, S8a and b). *E_0_’* showed ssDNA-like mechanical response (Fig. S8c) (30,44) similar to the mid-FRET population of hTel22 attributed to an unstructured ssDNA (Fig. S8c) (30). Like *E_1_* of hTel34, *E_1_’* unraveled gradually under tension (128 out of 186 events, ∼ 68 %, Figs. 5f and g). Like *E*_2_ of hTel_34_, *E_2_’* unfolded abruptly (58 events, Fig. 5f) at around two force clusters of ∼ 9 and 14 pN (Fig. S8e) while refolding occurred at lower forces (Figs. 5g, S8d and e). The gradual and reversible mechanical responses of the dominant populations of hTel34 (*E_1_*) or hTel34’ (*E_1_’*) can be likened with the non-cooperative unraveling via mutual strand slippage, observed with 3’ GQs in hTel28. This further hints at preferential GQ formation at the 3’ end in hTel34, similar to hTel28.

### Conformational dynamics of hTel28 and hTel34 in 100 mM Na^+^

We next investigated if the preferential GQ formation at the 3’ end of more than four telomeric repeats is dependent the cations present. Instead of hybrid GQ signatures in 100 mM K^+^, CD spectra of hTel28 in 100 mM Na^+^ showed signatures of antiparallel GQs (Fig. S9a) (45). In the absence of force, we observed a major population at *E* ∼ 0.72 (Fig. S9b). Under tension, unfolding predominantly occurred via an abrupt change in *E* from ∼ 0.72 to ∼ 0.1 at forces peaked at ∼ 8 pN (Figs. S10a, b and d) and refolding occurred at lower forces of ∼ 2.5 pN (Fig. S10e). Additionally, Type II partial unfolding was observed in ∼ 30 % (31 out of 104 events) of pulling cycles (Figs. S10a (cycle 3) and b). We did not observe Type I partial unfolding.

Among the hTel28 variants, stable GQs were formed only in hTel28.1, suggesting that GQ forms preferentially in the 3’ end also in Na^+^ solution (Fig. S9b). Other variants did not form stable GQs in Na^+^ likely because of poorer coordination by smaller Na^+^ compared to K^+^ (46).

Under tension, hTel28.1 predominantly showed Type II partial unfolding in 100 mM Na^+^ (87 out of 114 events, ∼ 76 %) (Figs. S10f and g) with the rest showing complete unfolding with two *f_unfold_* clusters centered at ∼ 5 and 19 pN (Figs. S10f-i). We next investigated hTel28.1T to test if an unpaired T_2_AG_3_ has any additional effect in Na^+^ that (dT)_6_ does not have. hTel28.1T showed a major population centered at *E* ∼ 0.75 in 100 mM Na^+^ (S11a). Under tension, it predominantly showed abrupt, complete unfolding with two *f_unfold_* clusters centered at ∼ 7 and 15 pN (Fig. S11b) with refolding occurring at lower forces (Fig. S11c). The rest (∼ 20 %, 22 out of 110 events) showed Type II partial unfolding. Overall, hTel28.1T was very similar to hTel28, both in its zero-force *E* value and its mechanical response. Thus, the GQ-core in Na^+^ remains unaffected by an additional T_2_AG_3_ overhang at the 3’ end.

hTel34 adopted anti-parallel GQ-like conformation in 100 mM Na^+^ and showed a single population centered at *E* ∼ 0.5 (Figs. S12a and b) (45). The mechanical response of hTel34 was gradual and reversible similar to *E_1_* in 100 mM K^+^ (Figs. S12c and d). However, this does not necessarily report on the dynamics of GQ within hTel34 because stretching of a single stranded overhang, if present, can overwhelm the response of the underlying GQ. To test the above possibility, we next interrogated hTel34’ behavior in 100 mM Na^+^ because it can report on the shorter, 22 nt, segment located in the 5’ end of hTel34. The major FRET population at *E* ∼ 0.72 (Fig. S12e) showed abrupt unfolding with three *f_unfold_* clusters of ∼ 4, 9 and 19 pN (Figs. S12h) and refolding at lower forces (Figs. S12f, g and i). Prior to the abrupt decreases in *E* we observed gradual decrease in *E* with increasing force, likely due to the presence of ssDNA region within folded hTel34’. We note that a 5’ GQ in hTel34’ would resemble hTel22 in FRET values with a main population at *E* ∼ 0.85 (30). Because hTel34’ shows a major conformation with *E* ∼ 0.72, we can eliminate the possibility of preferential GQ formation at the 5’ end in hTel34 (or hTel34’). A long-loop GQ in hTel34’, if it exists, would harbor a 15 nt long loop or two 9 nt long loops and have *E* ∼ 0.85 of hTel22 instead of the observed *E* ∼ 0.72 (30). Moreover, because long-loop GQs could not be stably formed in hTel28 in 100 mM Na^+^, we can rule out possible long-loop GQ formation in hTel34 and hTel34’. Thus, similar to 100 mM K^+^, our data are most consistent with GQ formation predominantly at the 3’ end of hTel34 in 100 mM Na^+^.

## Discussion

Despite its simple repetitive sequence, telomeric GQ possess extremely heterogeneous populations. Structural studies have revealed at least six different GQ conformations (e.g. parallel, antiparallel, hybrid etc.) based on strand symmetry, strand orientation and glycosidic conformation (18–24). Recently, we identified six mechanically different, interconvertible GQ species in a 22 nt long human telomeric DNA using single molecule fluorescence-force spectroscopy (30). In the context of longer telomeric sequences harboring more than four T_2_AG_3_ repeats, as we studied here using five repeats and six repeats using fluorescence-force spectroscopy, GQ polymorphism in tandem with their positional multiplicity along the DNA add to the complexity. Indeed, previous studies of four to seven repeats using smFRET only (hence at zero force) (32) or optical tweezers only (25) found evidence of multiple states. Using a series of mutants designed to mimic positionally defined GQs, we were able to separate positional multiplicity and GQ polymorphism.

### Five telomeric repeats

Here, we found that the mechanical response of GQ in a long telomeric DNA can depend on its location (Fig. 6a). For example, although both the 3’ GQ and the 5’ GQ in hTel28, mimicked by hTel28.1 and hTel28.5, respectively, utilize four consecutive T_2_AG_3_ segments, on application of force, the 3’ GQ former unravels gradually whereas the 5’ GQ withstands forces as high as 28 pN (Fig. S4c). This may stem from differences in the underlying GQ conformations as evidenced by their zero force *E* values (∼ 0.72 for hTel28.1 and ∼ 0.62 for hTel28.5, Fig. S4b). Because a GQ bearing a poly(dT) tail at the 5’ end had identical FRET value to a GQ with the same poly(dT) tail at the 3’ end (47), the different *E* values between hTel28.1 and hTel28.5 likely come from structurally different GQs (Figs. S4a and b). The gradual unfolding observed for 3’ GQ suggests a compliant structure, held together by local interactions which gradually get disrupted with the increasing force, reminiscent of mutual slippage where one guanine repeat slips relative to the other repeats one nucleotide at a time, disrupting individual G-quartets one at a time (48). Such a mode of unraveling is conceivable for parallel GQs as slippage of a G-strand does not perturb the inherent all *anti-*G configuration of the remaining G-quartets. However, as antiparallel or hybrid GQs are each characterized by a unique arrangement of *anti* and *syn*-Gs, strand slippage upsets the inherent G arrangement in a quartet and disarrays the entire structure cooperatively.

**Fig. 6:**
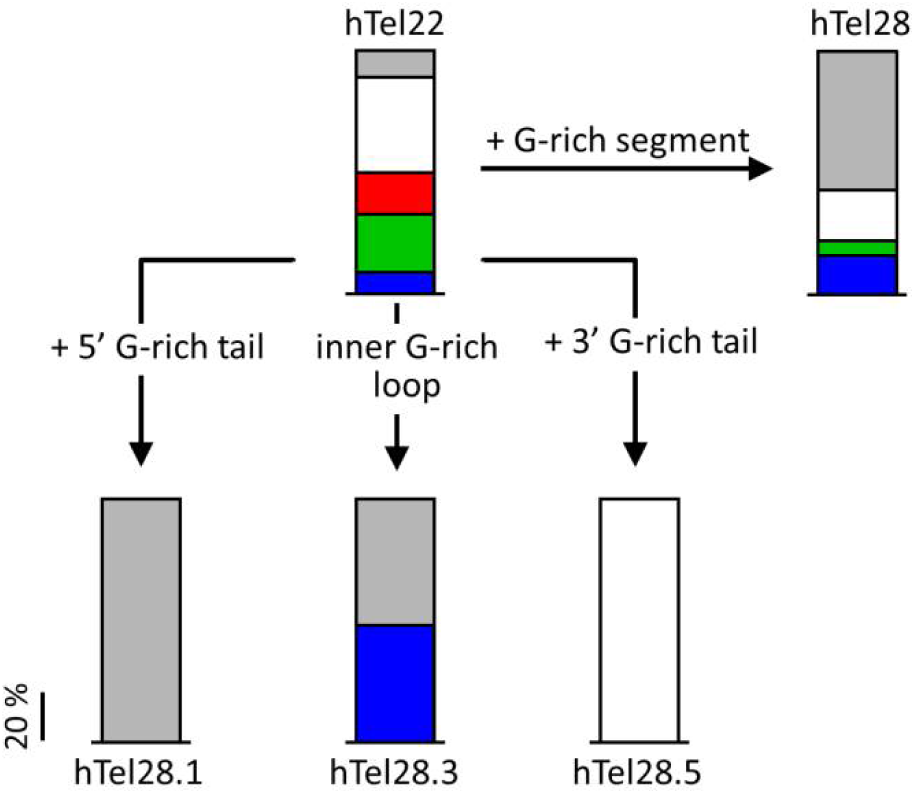
Effect of an unpaired G-rich segment in canonical GQs (hTel22) in 100 mM K^+^. The mechanical states are differentiated based on *Δx*^‡^s of ∼ 6.8 (blue), 2.8 (green) and 1.8 nm (red) for complete unfolding. Grey and white bars represent partially unfolded and ultrastable states respectively. Mechanical response of hTel28, which is hTel22 with a G-rich segment, is a cumulative effect of hTel28.1, hTel28.5, hTel28.3 etc.

Among the variants of hTel28, only a long-loop variant, hTel28.3, showed cooperative destabilization of the G-quartets, marked by abrupt unfolding under increasing force. Analysis of the corresponding *f_unfold_* histogram, which is peaked at 7.5 pN, yielded free energy parameters *Δx*^‡^ (distance to the transition state) and *ΔG*^‡^s (apparent free energy of activation) as ∼ 6.6 nm and ∼ 7 k_B_T, respectively (Fig. S13a). In contrast, complete unfolding of hTel28 showed two *f_unfold_* clusters at ∼ 7.5 and 17 pN, corresponding to *Δx*^‡^s of ∼ 6.7 and 3.2 nm, respectively (Fig. S13a). A *Δx*^‡^ of ∼ 6.1 nm was estimated for a mechanically weak subpopulation (*f_unfold_* < 5 pN) of hTel22 under identical buffer conditions (30), attributed to partially folded GQs containing two G-quartets instead of three (30). Because *Δx*^‡^ of the 7.5 pN cluster of hTel28 and hTel28.3 is close to that of the two-tiered hTel22-GQs, we suggest the presence of frayed G-quartet structures adjacent to the inner T_2_AG_3_-containing loop. The 17 pN cluster can be attributed to hTel28.2 or hTel28.4 because it was not seen for hTel28.1, hTel28.3 or hTel28.5.

Although our single molecule studies suggest preferential GQ formation at the 3’ end of hTel28, such GQs with a 5’ T_2_AG_3_ overhang were mechanically distinct from GQs formed in hTel22 (Fig. 6). Partial and gradual unfolding attributed to mutual slippage, which was a minority behavior in hTel22, dominates the overall mechanical response of hTel28 (30). Interestingly, upon substitution of the hexanucleotide G-rich strand with a poly(dT) segment of the same length (hTel28.1T), the canonical hTel22 like behavior was restored. In addition to ultrastable (∼ 35 %) and partially unfolded GQs (∼ 10 %), we observed at least three cooperatively unfolding species with *f_unfold_*s of ∼ 3.5, 10 and 20 pN from hTel28.1T. The respective *Δx*^‡^s were estimated as ∼ 6.8, 2.8 and 1.8 nm (Figs. 6 and S13a), close to the previously reported values of ∼ 6.1, 3.1 and 1.8 nm in hTel22 (30).

In all, our study highlights the regulatory role of an unassociated G-rich strand on the GQ core of a five-repeat telomeric sequence. A T_2_AG_3_ overhang at the 5’ end biases the molecule toward the partial unfolding pathway whereas when present at the 3’ end it renders GQ ultrastable. On the other hand, an inner loop harboring a T_2_AG_3_ segment manifests cooperative unfolding similar to canonical hTel22-GQs (30). We hypothesize position-dependent transient interactions between the unpaired T_2_AG_3_ segment with the underlying GQ core, which directly affects the conformational distribution by streamlining a mechanically diverse population to a single conformation (Fig. 6). Interactions between unpaired G-nucleotides and the GQ core was also observed in the GQ-forming c*-Myc* promoter (49). However, detailed structural studies are required to understand the exact nature of such interactions. Nonetheless, at applied forces of up to ∼ 28 pN, because the 3’ GQs in hTel28 are only partially unraveled, they rarely convert into long-loop GQs (∼ 16 %) upon relaxation. In contrast, a long-loop GQ, when unfolded cooperatively, loses memory of its initial state and can relax into a mechanically different long-loop GQ or even a 3’ GQ (∼ 33 %).

### Six telomeric repeats

From hTel34 which contains six human telomeric repeats, we observed two major populations *E*_1_ and *E*_2_ at zero force. To reduce the compounding effect of stretching of ssDNA tail(s), we also designed the internally labeled construct hTel34’ whose FRET response is sensitive to the conformation of the four 5’ most repeats. By comparative analysis of responses of hTel28 and hTel34, we assigned the major population, *E_1_* (or *E_1_’*) of hTel34 (or hTel34’) to 3’GQ with a 12 nt ssDNA overhang harboring two T_2_AG_3_ segments. The minor populations, *E_2_* (hTel34) and *E_2_’* (hTel34’) show abrupt, cooperative unfolding under tension similar to long-loop GQs in hTel28. The *f_unfold_* of *E_2_* reveals two force clusters at ∼ 8 and 16 pN with underlying *Δx*^‡^s of ∼ 3.7 and 4 nm respectively (Fig. S13b). Although similar in *Δx*^‡^, the cluster at *f_unfold_*=16 pN had significantly higher *τ_u_*(0) and *ΔG*^‡^ (10990 vs 59.83 s and 11.6 vs 5.7 k_B_T), indicating presence of long and short-lived species that are structurally similar (24). The *f_unfold_* clusters in *E_2_’* correspond to *Δx*^‡^s of ∼ 3.6 and 3.7 nm respectively, similar to *E_2_*. Hence, we suggest that *E_2_* of hTel34 are *E_2_’* of hTel34’ represent the same conformation (Fig. S13b).

GQs formed at the 5’ end of hTel28 were ultrastable. Analogous behavior might be overshadowed by ssDNA stretching in hTel34, but should be reflected as stable high FRET over the complete pulling cycle in hTel34’. Although *E_2_’* showed a no-force FRET value of ∼ 0.88, expected for a 5’ GQ, it did not show stable high FRET under tension. Rather, *E_2_*’ showed abrupt unfolding, reminiscent of long-loop GQs in hTel28. So we suggest that *E*_2_ of hTel34 is unlikely due to a 5’ GQ. Can it be due to a 3’ GQ? A 3’ GQ would leave a 12 nt ssDNA region in hTel34’ which can be stretched under tension, giving rise to initial gradual FRET decrease. Because we did not observe any FRET decrease until abrupt unfolding, we can also rule out the possibility that *E*_2_’ and *E_2_* correspond to a 3’ GQ. Overall, our data suggest that *E_2_* represents long-loop populations although we cannot tell where there is a single long loop of 15 nt in length or two separate 9 nt long loops. The difference in transition distances between hTel34 and hTel28 may be attributed to structural differences stemming from the size and arrangement of the intervening loop(s).

### Dynamics in Na^+^ solution

In 100 mM Na^+^, hTel28 showed the abrupt unfolding response in majority of events, reminiscent of hTel22. We estimated *Δx*^‡^ *τ_u_*(0) and *ΔG*^‡^ as ∼ 4.3 nm, ∼ 50 s and 6.5 k_B_T, respectively. Among the variants of hTel28, only hTel28.1 showed stable folding in Na^+^, suggesting that 3’ GQ formation is favored also in Na^+^. Upon substituting the hexanucleotide G-rich segment at the 5’ end with (dT)_6_, we observed two *f_unfold_* clusterscorresponding to long-lived (*τ_u_*(0) ∼ 1190 s, *f_unfold_* ∼14.8 pN) and short-lived (*τ_u_*(0) ∼ 68 s, *f_unfold_* ∼6.8 pN) isostructural species with *Δx*^‡^ of ∼4 nm, which is close to a previously reported value for hTel22 (Fig. S15) (30). Thus, GQ conformation is not significantly affected by the presence of a T_2_AG_3_ overhang at the 5’ end in 100 mM Na^+^. We estimated similar *Δx*^‡^s in 3’ GQ forming hTel28 and hTel34’ (Fig. S14). This substantiates our hypothesis that GQ forms preferentially at the 3’ end of hTel34 also in Na^+^.

Taken together, our study suggests a complex, sequence dependent interplay between GQ and the unpaired Gs in five and six-repeat telomeric DNA. We note that, the predominant Type I partial unfolding observed in hTel28 and hTel34 is exclusive to 100 mM K^+^. Thus, although GQ formation is polarized at the 3’ end in Na^+^ and K^+^, the streamlining effect of an associated telomeric repeat on GQ folding appears unique to K+ conditions.

### Concluding remarks

G-rich telomeric sequences are well known to form stable GQs *in vitro*. In cells, telomeric DNA spans over 100-200 nt, beyond the double-stranded region, in the form of multiple G-rich hexanucleotide repeats. Using single molecule fluorescence-force spectroscopy, we demonstrated preferential GQ formation at the 3’ end of telomeres under physiologically relevant conditions. Although the sequences used in this study can harbor only one GQ, our results might imply a general scheme of polarized GQ formation being initiated from the 3’ end of long telomeric DNA, spanning multiple repeats of GQ-forming segments. The relative position of GQ(s) with respect to the 3’ end, i.e. the size of ssDNA tail downstream of the 3’ end, has important implications in processes such as extension by telomerase and alternate lengthening of telomeres (ALT). Wang *et al* showed that a minimum 3’ ssDNA tail of approximately 6 nt, 8 nt and 12 nt in length is essential for helicase unwinding of telomeric GQ, extension by telomerase, and lengthening by ALT, respectively (14,50–52). Thus GQ formation at the 3’ end of the telomeric DNA is likely to inhibit the aforementioned processes. On the other hand, although a population minority, the long-loop GQ species might be more susceptible to helicase-catalyzed by providing an internal loading site for helicases (53). Telomeres are assumed to be protected by T-loop formation, which involves invasion into the double-stranded telomeric region by the G-rich telomeric overhang and subsequent annealing with the C-rich strand (54). Stable GQs at the 3’ end might interfere with T-loop formation as well. In the cellular milieu, processes such as transcription and replication generate tension. However, our study shows that long telomeric repeats cannot be fully unwound at average stall forces of motor proteins and polymerases (55,56). Thus, GQs can act as potential kinetic traps and inhibit replication and transcription.

## Acknowledgements

We thank Momčilo Gavrilov for providing Rep-X. This work was supported by National Science Foundation Grant PHY-1430124 (to T.H.) and National Institutes of Health Grants GM122569 (to T.H.). T.H. is an Investigator with the Howard Hughes Medical Institute.

