## Supporting Information for "Streamlining effects of extra telomeric repeat on telomere folding revealed by fluorescence-force spectroscopy"

*Jaba Mitra[1,2] and Taekjip Ha[2,3,4,5*]*

*[1]Department of Materials Science and Engineering, University of Illinois at Urbana-Champaign, Urbana IL, USA; [2]Department of Biophysics and Biophysical Chemistry, Johns Hopkins University, Baltimore, MD, USA; [3]Department of Biophysics, Johns Hopkins University, Baltimore, MD, USA; [4]Department of Biomedical Engineering, Johns Hopkins University, Baltimore, MD, USA; [5]Howard Hughes Medical Institute, USA*

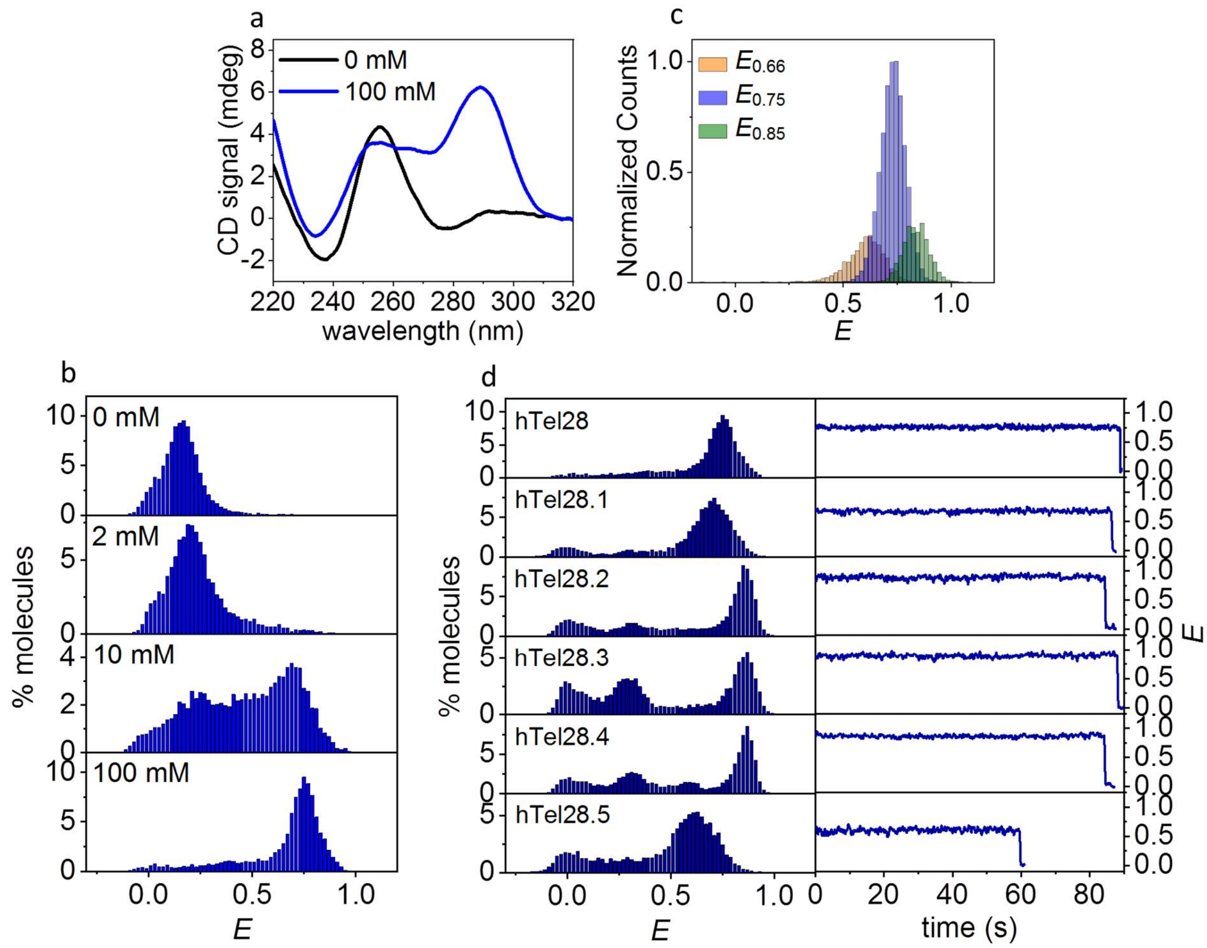


**Fig. S1**: (a) CD spectra of hTel28 in 0 and 100 mM $K^+$. (b) *E* histograms of hTel28 as a function of $K^+$ concentration. (c) *E* histograms recast from time trajectories of hTel28 in 100 mM $K^+$, at *E* values of ~ 0.68, 0.75 and 0.86. (d) *E* histograms (left) and single molecule time trajectories (30 ms integration time, right) of hTel28 mutants in 100 mM $K^+$ in the absence of force.

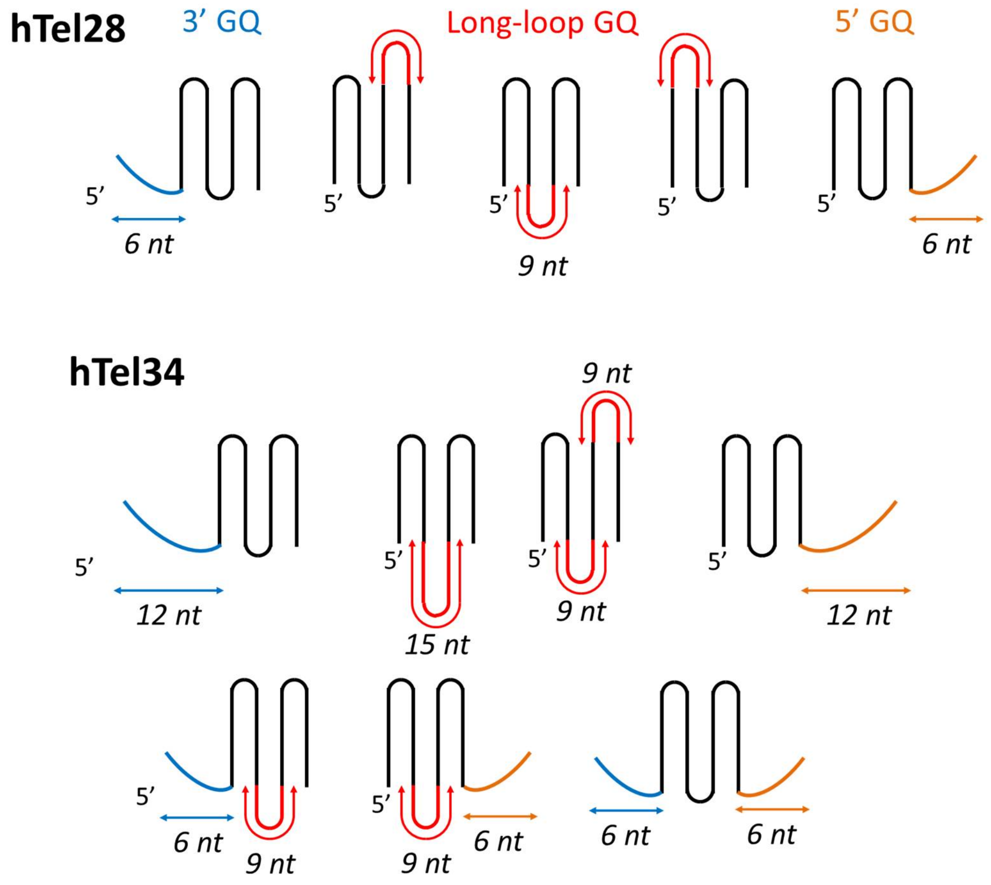


**Fig. S2**: GQ formation in hTel28 and hTel34. In hTel28, GQs can form at the termini i.e. 3’ or 5’ end (hTel28.1 and hTel28.5 respectively) or by association of a 9 nt long loop bearing a $T_2AG_3$ segment (e.g. hTel28.2, hTel28.3 or hTel28.4). The unassociated $T_2AG_3$ segment at the 5’ and 3’ ends are shown in blue and orange respectively whereas the long loop is shown in red. The conformations represent hTel28.1 to hTel28.5 from left to right.

In hTel34, GQ can form at the 3’ or 5’ end with a 12 nt long ssDNA tail (two conformations differing in GQ polarity). Long-loop GQs can harbor a 15 nt long loop harboring two $T_2AG_3$ segments (3 conformations), two 9 nt long loops with a $T_2AG_3$ segment each (3 conformations) or a 9 nt loop with a ssDNA tail at the 5’ or 3’ end (6 conformations). Additionally, GQ formation may involve two 6 nt overhangs at both the 3’ and 5’ ends (1 conformation). Fifteen conformations in all can be conceived based on the above prototypes.

The conformations illustrated only take into account the differences in the relative position of the unpaired $T_2AG_3$ segment, irrespective of the structure of the GQ formed.

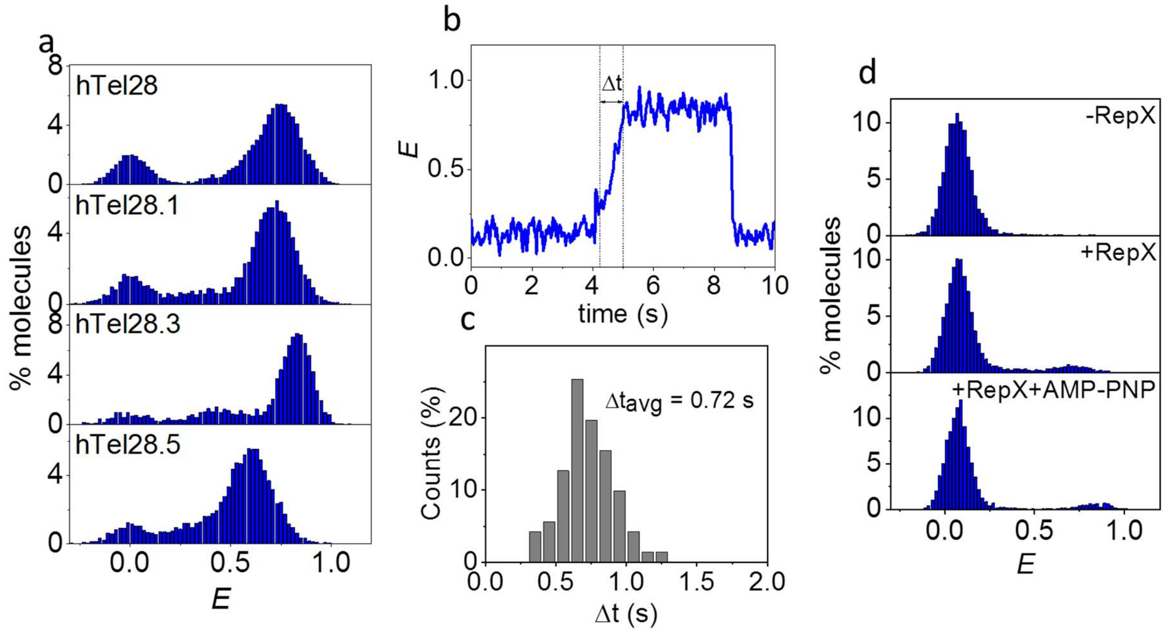


**Fig. S3**: (a) *E* histograms of hTel28, hTel28.1, hTel28.3 and hTel28.5 in the Rep-X unwinding buffer. (b) Real-time *E* trajectory of unwinding of hTel28-duplex and folding of GQ in Δt time (30 ms integration time). (c) Histogram of Δt. (d) *E* histograms of hTel28-duplex in the Rep-X unbound (top) and bound (middle) states, and after Rep-X unwinding of the duplex in presence of 1 mM AMP-PNP and 2.5 mM $MgCl_2$ (bottom).

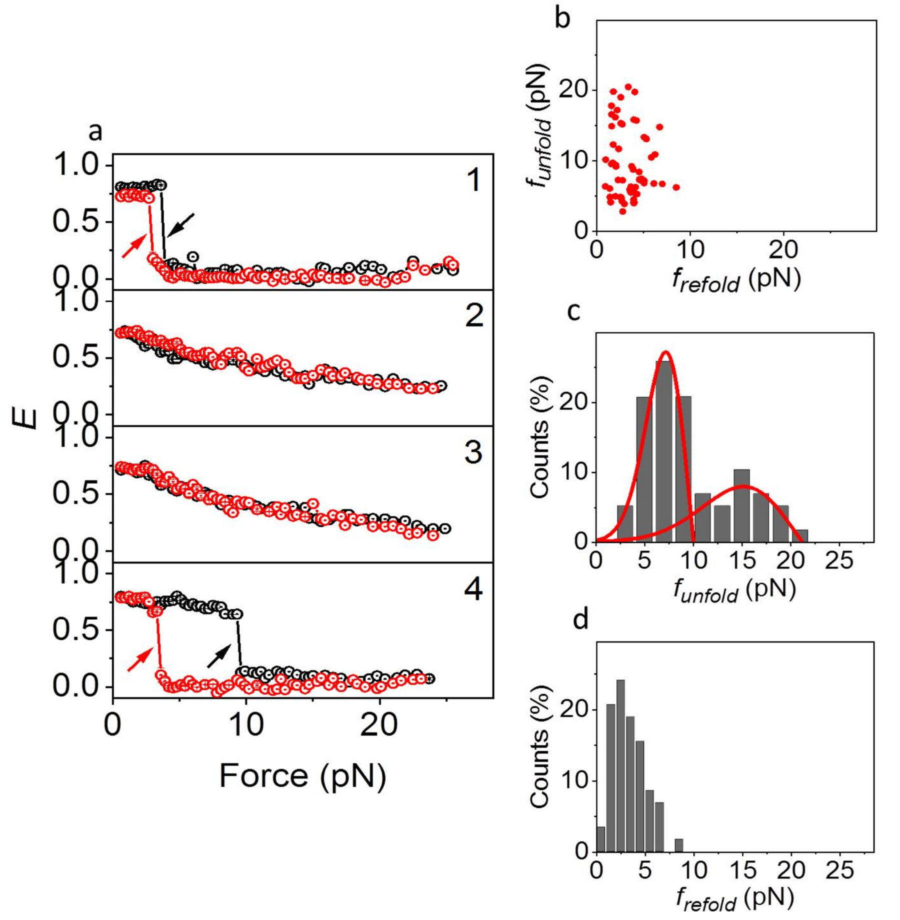


**Fig. S4**: (a) *E* vs force response of the hTel28 molecule shown in Fig. 3b. The pulling cycle # is depicted in the inset. Stretching and relaxation are represented in black and red respectively. Cycles 1 and 4 show complete unfolding, whereas cycles 2 and 3 show partial (I) unfolding. The arrows denote $f_{unfold}$ (black) and $f_{refold}$ (red). (b) A scatter plot of $f_{unfold}$ and $f_{refold}$ of complete unfolding events (*N*=58). (c and d) $f_{unfold}$ (c) and $f_{refold}$ (d) histograms corresponding to complete unfolding events (*N*=58). The red curves represent the rupture force distributions predicted using the Dudko-Szabo model (1,2). All measurements were done in 100 mM $K^+$.

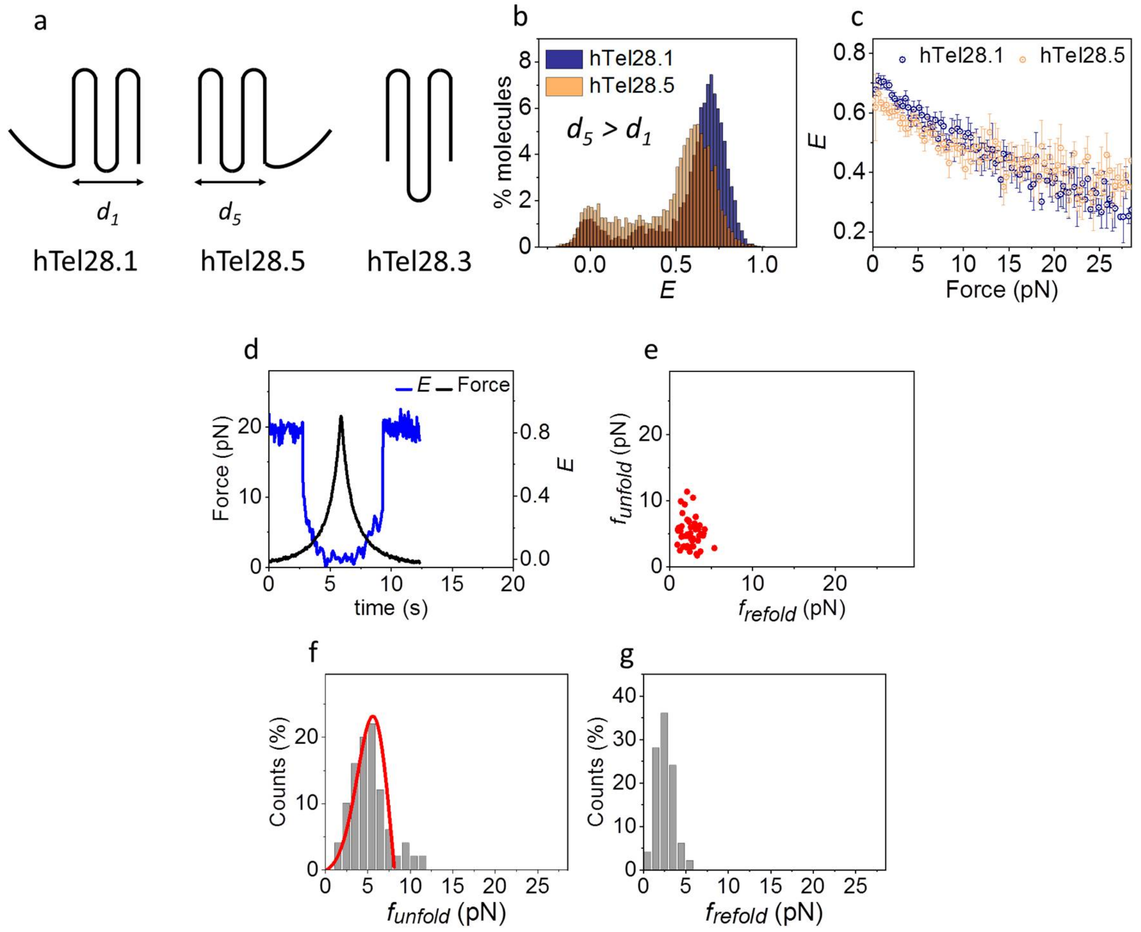


**Fig. S5**: (a) A cartoon depicting polarized GQ formation at the 3' (hTel28.1) and 5' (hTel28.5) end or via association of a long loop (hTel28.3). The end to end distances of GQs formed in hTel28.1 and hTel28.5 (excluding the ssDNA tail), are represented by $d_1$ and $d_5$ respectively. (b) Overlap of *E* histograms of hTel28.1 and hTel28.5 in 100 mM $K^+$. (c) Overlap of average *E* vs force response (stretch) of hTel28.1 and hTel28.5 in 100 mM $K^+$. (d) A representative *E* time trace of hTel28.3 in one pulling cycle (20 ms integration time). (e) A scatter plot of $f_{unfold}$ and $f_{refold}$ ($N$=50). (f and g) $f_{unfold}$ (f) and $f_{refold}$ (g) histograms corresponding to complete unfolding events ($N$=50). The red curve represents the rupture force distribution predicted using the Dudko-Szabo model (1,2). All measurements were done in 100 mM $K^+$.

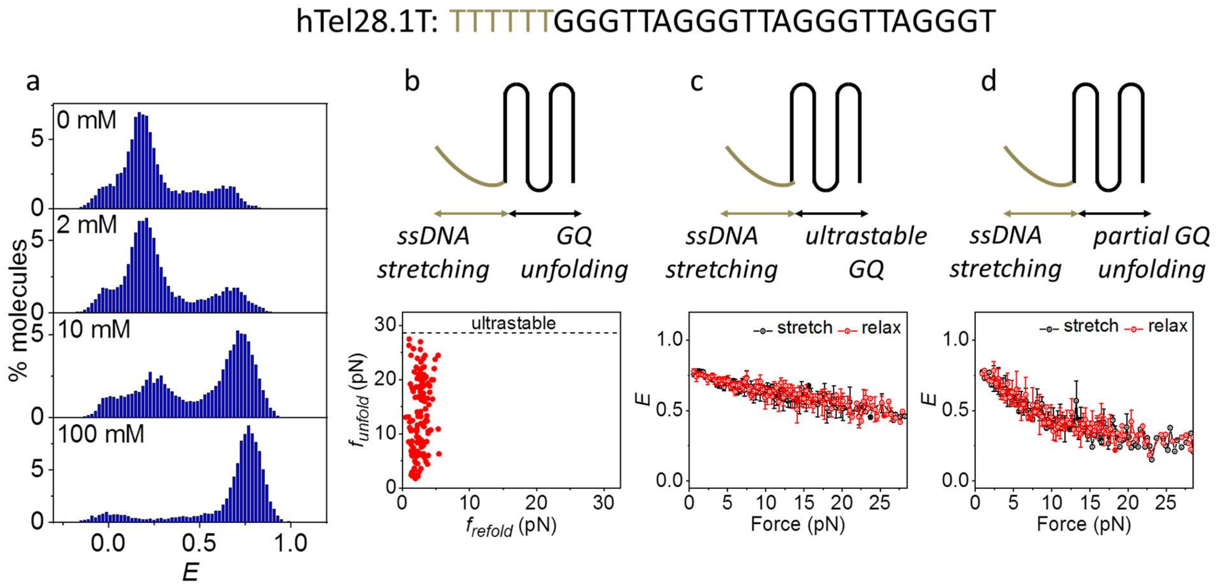


**Fig. S6**: (a) $E$ histograms of hTel28.T as a function of $K^+$ concentration. (b-d) Schematic depicting three different mechanical responses of hTel28.1T in 100 mM $K^+$. A scatter plot of $f_{unfold}$ and $f_{refold}$ is shown in (b, lower panel, $N$=131). Average $E$ vs force responses of ultrastable and partially unfolded GQs are shown in (c, $N$=84) and (d, $N$=24).

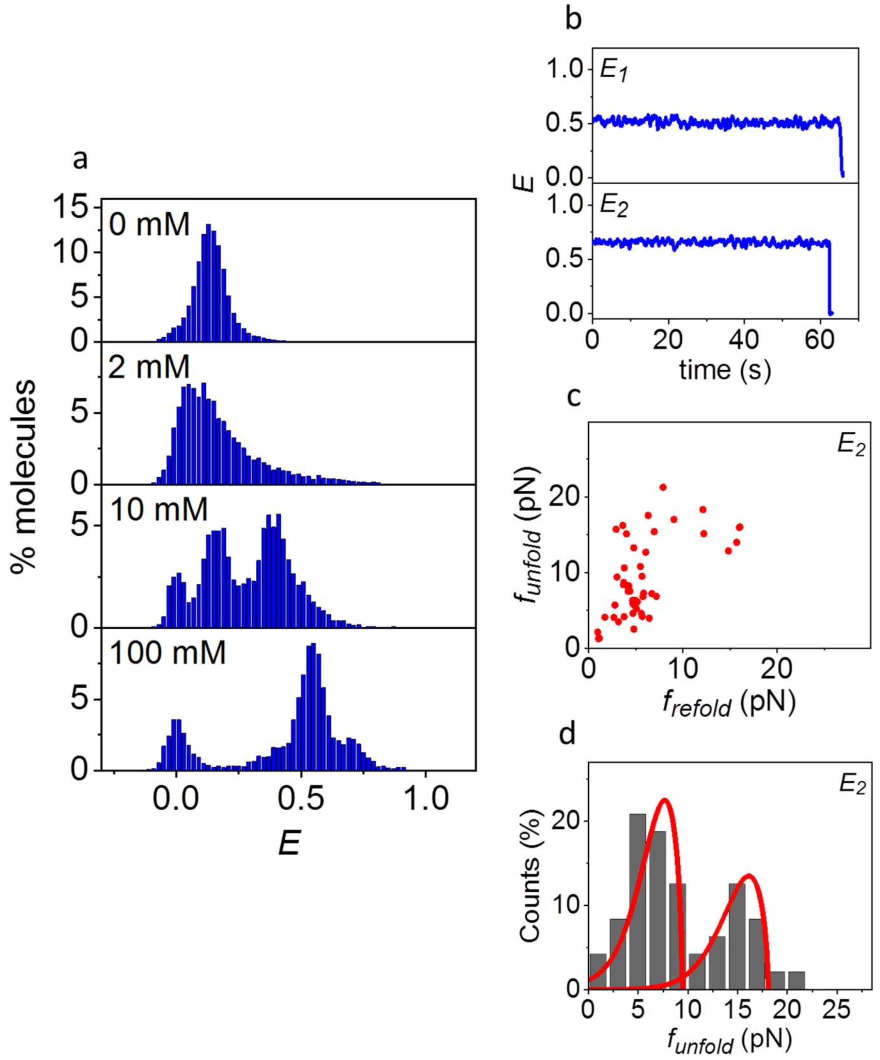


**Fig. S7**: (a) *E* histogram of hTel34 as a function of $K^+$ concentration in the absence of force. (b) Representative *E* time traces of $E_1$ and $E_2$ (30 ms integration time) in absence of force. (c) A scatter plot of $f_{unfold}$ and $f_{refold}$ of $E_2$. (d) Histogram of $f_{unfold}$. The red curves represent the rupture force distributions predicted using the Dudko-Szabo model (1,2). *N*=48 for (c) and (d).

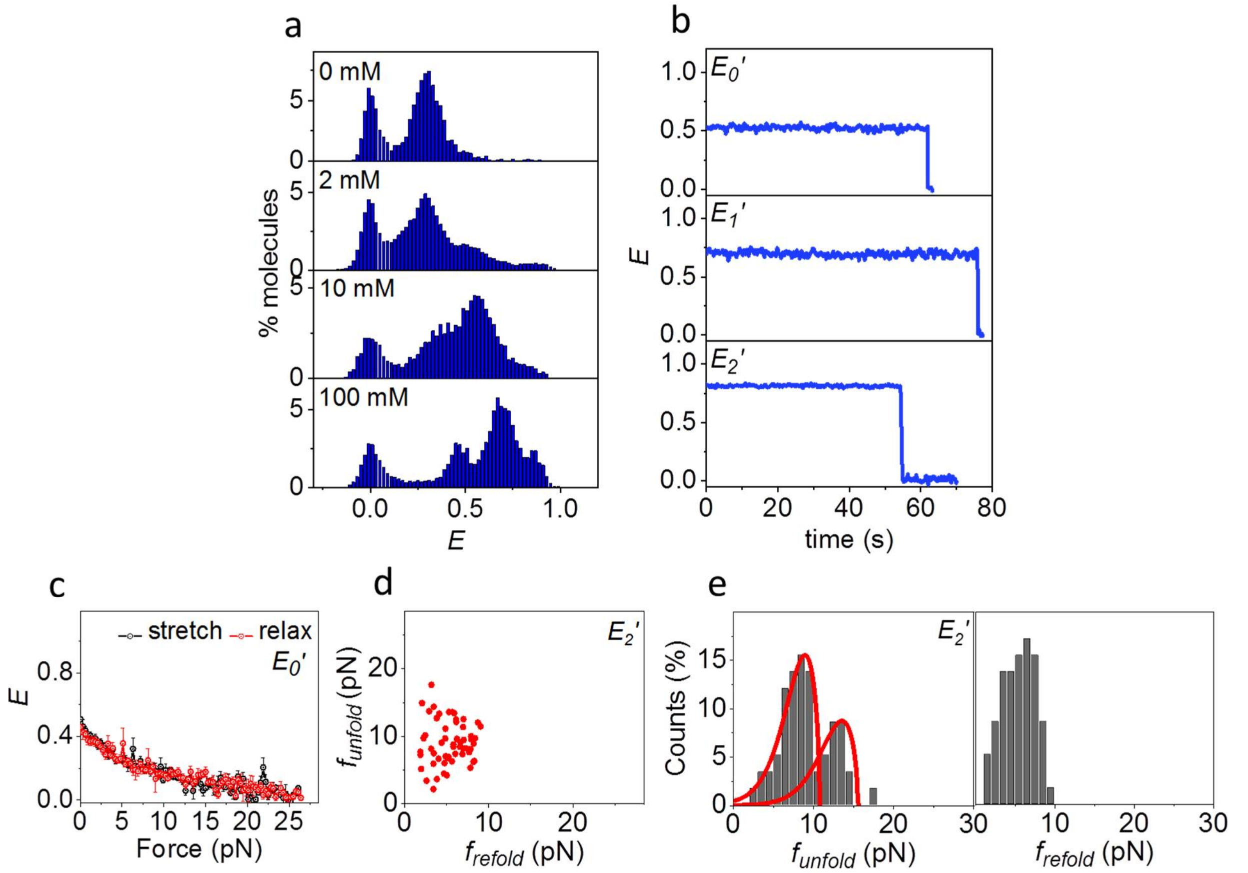


**Fig. S8**: (a) $E$ histogram of hTel34' as a function of $K^+$ concentration in the absence of force. (b) Representative $E$ time traces of $E_0$', $E_1$' and $E_2$' in 100 mM $K^+$ in absence of force (30 ms integration time). (c) Average $E$ vs force response of $E_0$'. (d) A scatter plot of $f_{unfold}$ and $f_{refold}$ of $E_1$'. (e) Histograms of $f_{unfold}$ and $f_{refold}$ for $E_1$'. The red curves represent the rupture force distributions predicted using the Dudko-Szabo model (1,2). $N$=58 for (d) and (e).

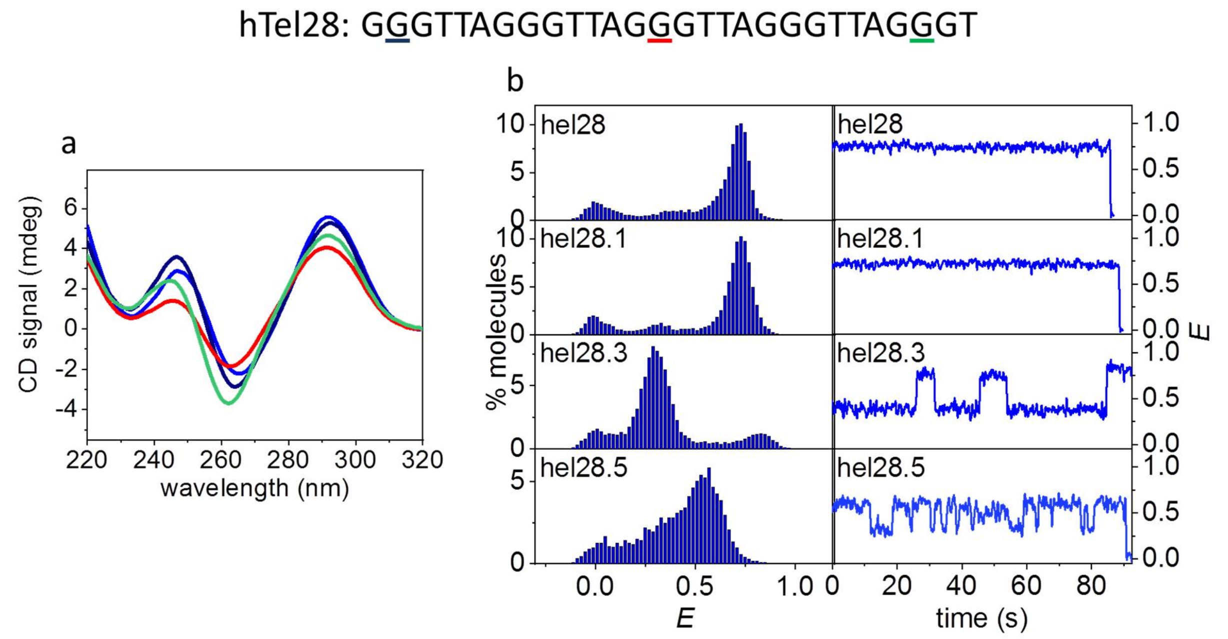


**Fig. S9**: (a) CD spectra of hTel28 (blue), hTel28.1 (navy blue), hTel28.3 (red) and hTel28.5 (green) in 100 mM $Na^+$. The G to T mutation sites in hTel28 corresponding to hTel28.1, hTel28.3 and hTel28.5 are underlined on the sequence with navy blue, red and green respectively. (b) *E* histograms (left) and representative time traces (right, 30 ms integration time) of hTel28 variants in 100 mM $Na^+$ in absence of force.

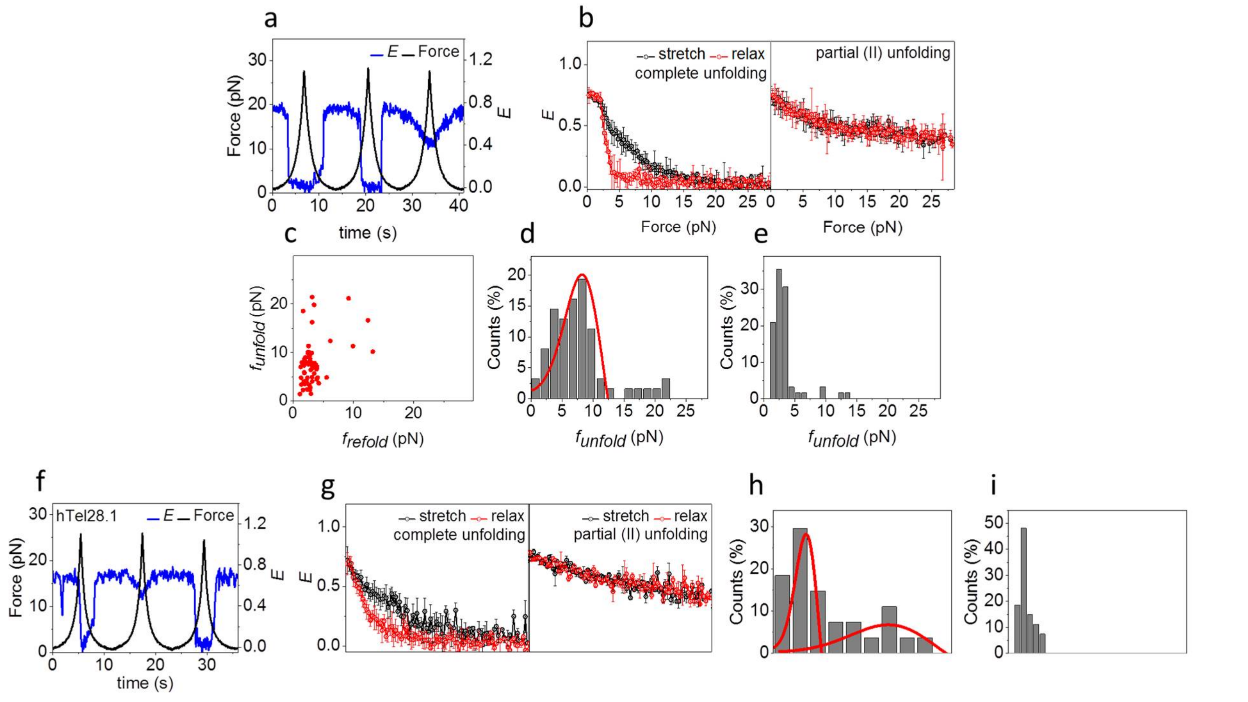


**Fig. S10**: (a) A representative *E* time trace of hTel28 in three pulling cycles. (b) Average *E* vs force response of molecule showing complete (*N*=73, left) and partial (II) (*N*=31, right) unfolding. (c) A scatter plot of $f_{unfold}$ and $f_{refold}$ corresponding to (b, left) ($N$=73). (d and e) Histograms of $f_{unfold}$ (d) and $f_{refold}$ (e) corresponding to complete unfolding events only. (f) Representative $E$ time trace of hTel28.1. (g) Average $E$ vs force response of hTel28.1. (h and i) Histograms of $f_{unfold}$ (h) hTel28.1 and $f_{refold}$ (i) of hTel28.1 showing complete unfolding events only. ($N$=27 for hTel28.1). All measurements were done in 100 mM $Na^{+}$, at an integration time of 20 ms.

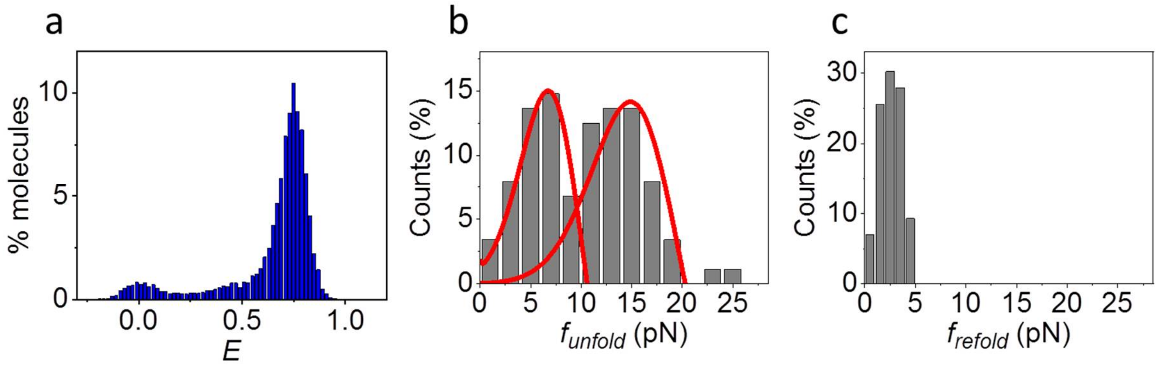


**Fig. S11**: (a) $E$ histogram of hTel28.1T in 100 mM $Na^{+}$, in the absence of force. (b and c) Distributions of $f_{unfold}$ (b) and $f_{refold}$ (c) ($N$=88). The red curves represent the rupture force distributions predicted using the Dudko-Szabo model (1,2).

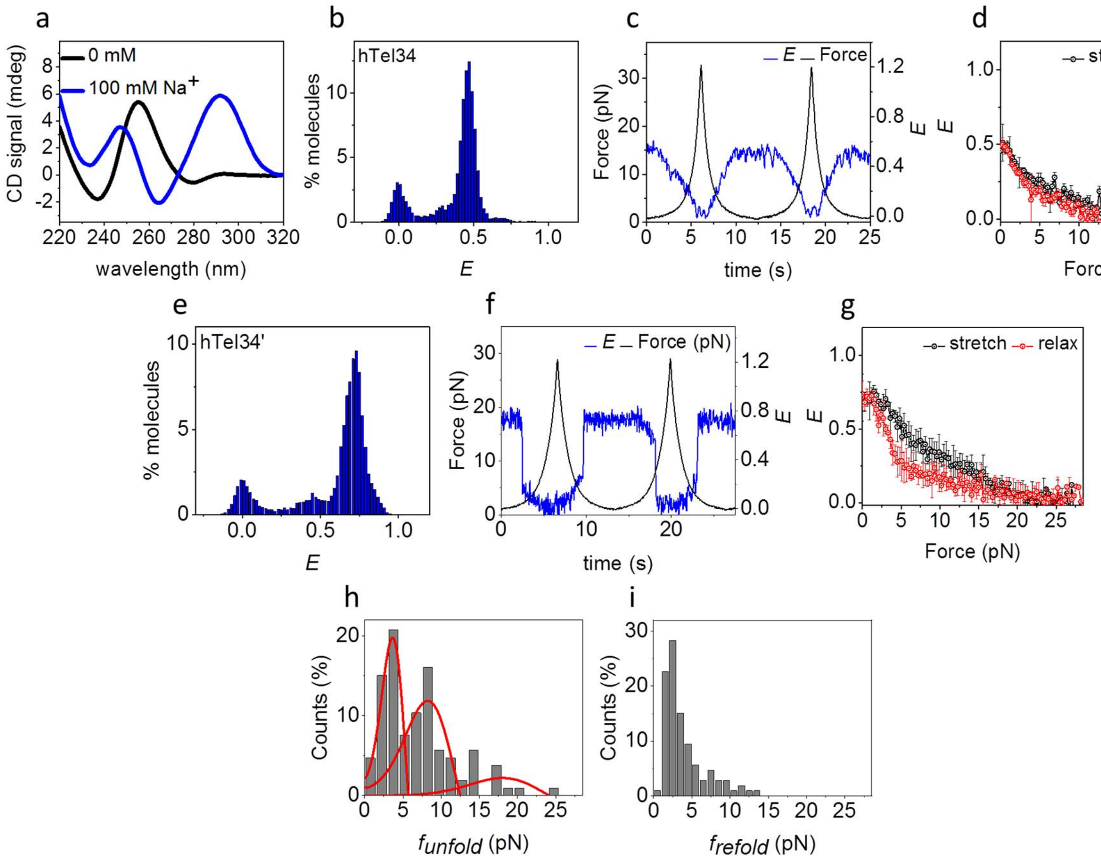


**Fig. S12**: (a) CD spectra of hTel34. (b) $E$ histogram of hTel34 in the absence of force (c) A representative $E$ time trace showing two pulling cycles. (d) Average $E$ vs force response of hTel34. (e) $E$ histogram of hTel34’. (f) A representative $E$ time trace showing three pulling cycles. (g) Average $E$ vs force response of hTel34’. (h and i) Histograms of $f_{unfold}$ (h) and $f_{refold}$ (i) ($N$=105). The red curves represent the rupture force distributions predicted using the Dudko-Szabo model (1,2). The data in (b-i) were collected in 100 mM $Na^+$. (c) and (f) were collected at an integration time of 20 ms.

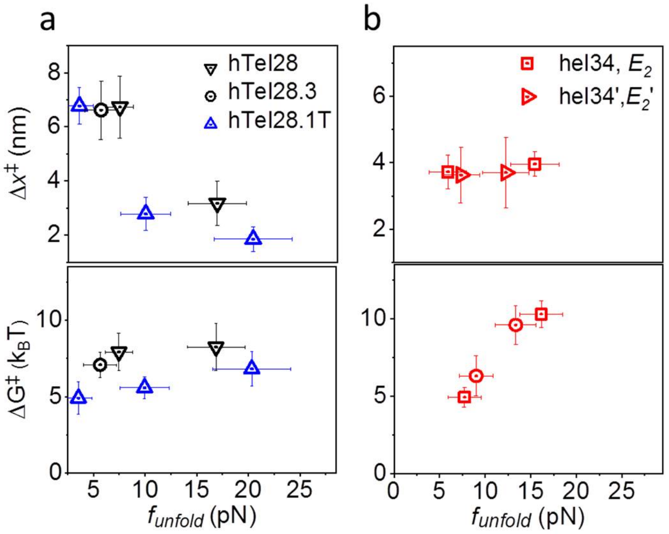


**Fig. S13**: $\Delta x^{\ddagger}$s (top) and $\Delta G^{\ddagger}$s (bottom) corresponding to the force clusters observed in hTel28, hTel28.3 and hTel28.1T (a) and hTel34 and hTel34’ (b) in 100 mM $K^+$.

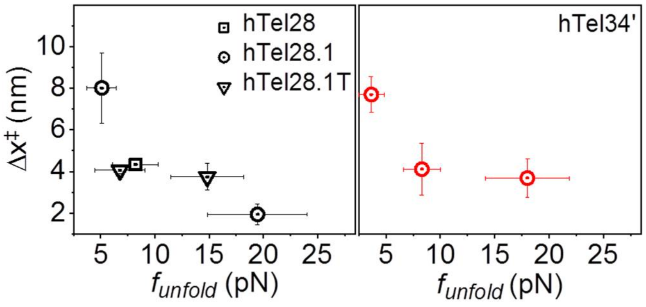


**Fig. S14**: $\Delta x^{\ddagger}$s corresponding to the force clusters observed in hTel28, hTel28.1, hTel28.5, hTel28.1T and hTel34’ in 100 mM $Na^+$.

All error bars represent standard errors.
